# Ultrafast elastocapillary fans control agile maneuvering in ripple bugs and robots

**DOI:** 10.1101/2024.11.15.623609

**Authors:** Victor M. Ortega-Jimenez, Dongjin Kim, Sunny Kumar, Changhwan Kim, Je-Sung Koh, Saad Bhamla

## Abstract

Millimeter-sized ripple bugs in the genus *Rhagovelia* exhibit exceptional agility and rapid maneuvers in fast, unsteady streams, comparable to animal fliers. Their remarkable interfacial transit and turning skills stem from a specialized fan structure on their middle legs. While researchers have suggested active fan actuation, the role of capillary forces and unique microstructure in self-spreading remains unclear. We reveal that *Rhagovelia’s* fans possess a flat-ribbon architecture with directional stiffness, enabling ultrafast elastocapillary morphing for passive actuation in under 10 ms, independent of muscle control, while producing high-thrust momentum through unsteady vortical wakes. These self-morphing fans allow *Rhagovelia* to execute ∼90 ° turns at a rate of ∼4200 °/s, in ∼50 ms, with speeds reaching ∼120 BL/s – on par with the fastest recorded turns in animal fliers like fruit flies. Inspired by these, we develop an ultralight, ultrafast elastocapillary robotic fan (∼1 mg, ∼100 ms opening/closing time) and integrate it into an insect-scaled robot (Rhagobot, ∼0.2 g). The engineered fans passively balance surface tension and water drag through stiffness anisotropy, enabling the robot to achieve high agility with speeds up to ∼2 BL/s and turning rates of 206 °/sec. Experiments with both insects and robots, with and without fans, show that a self-spreading passive fan significantly improves thrust, braking, and turning – key factors for controlled, high-speed maneuvers. This elastocapillary innovation enables ripple bugs to survive and thrive in turbulent streams and offers new insights for agile aquatic robotics.

---

Ripple bugs (Veliidae: Rhagovelia) exhibit remarkable diversity around the globe^1-3^,and are distinguished by their millimeter size and agility in navigating turbulent streams and coastal waters. This locomotory proficiency stems from a unique adaptation: a feathery fan-like structure at the distal end of their middle legs (Fig. 1a). Recent research shows that this fan, regulated by two taxon-restricted genes^4^, enhances maneuverability in an ecologically challenging niche by increasing turning angles and facilitating movement upstream in rapid flows^4^. The fan’s feather-like microstructure, which promotes wetting rather than water repellency, presents an intriguing biomechanical puzzle: how do the fan’s geometry and elastocapillarity drive the rapid and reversible self-immersion, opening, and closure of the fan’s barbs and barbule?

**Fig. 1.**
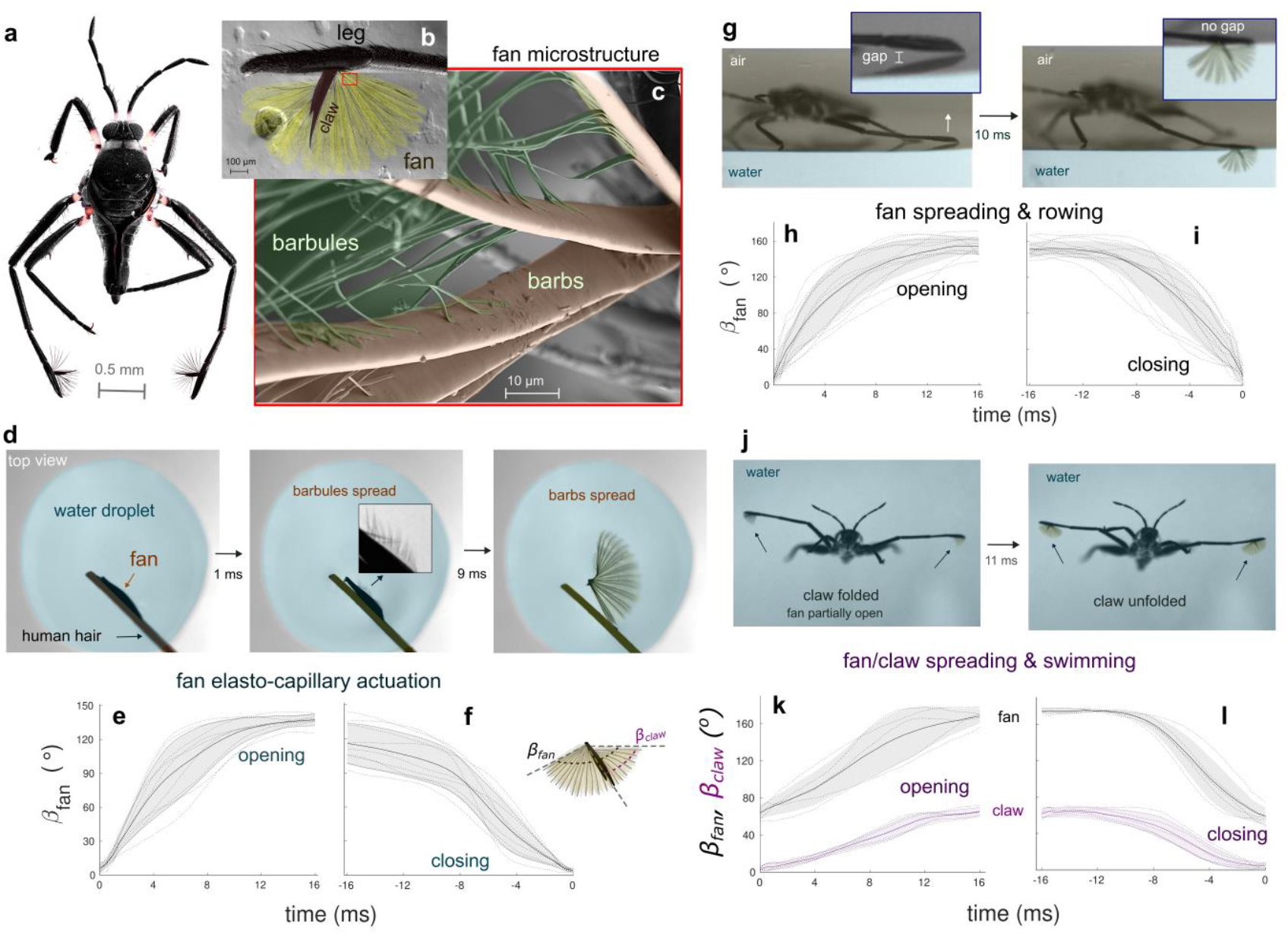
Rhagovelia fan microstructure and spreading performance. **a**, *Rhagovelia obesa* showing fans and claws at the tip of the middle legs. **b**, Image of fan and claw. **c**, SEM image of flat-ribbon microstructure of barbs and barbules. **d**, Video frames showing a top view of an isolated fan before (left) and while placed on a water droplet. Barbules spread after 1 second (middle) and barbs after 10 seconds (right). Time series of the spreading angle (*β*_*fan*_) of the isolated fan during opening **(e)** and closing **(f). g**, Video frames of a rowing individual before (left) and when the fan is fully open in the water (right). Time series of fan’s spreading angle (*β*_*fan*_) during opening **(h)** and closing **(i). j**, Video frames of a swimming individual before (left) and when the fan is fully open (right). The fan remains partially spread before the claw starts to open. Time series of the fan’s and claw’s spreading angle (*β*_*fan*_ and *β*_*claw*_) during opening **(k)** and closing **(l)**. Lines, shadows, and dashed lines in plots represent the mean value, one s.d., and individual data, respectively.

Rhagovelia’s feathered fan provides an inspirational template for developing self-morphing artificial propellers, offering insights into their biological form and function. Implementing this biological innovation in artificial systems could improve the maneuverability of interfacial microrobots. However, such configurations are largely unexplored in semi-aquatic robots^5,6^. Current semi-aquatic robots, designed for moving on still water surfaces, utilize three primary size-dependent propulsion mechanisms based on their size^5^. Large-scale robots (>50 g) row using bulky, rigid hydrophilic pads (mimicking oars) to generate drag during leg strokes^7-11^, while small-scale robots (< 50g) row with thin, hydrophobic legs to minimize surface tension, ^12-19^ mimicking Geriidae water striders^20^. At small scales, surface tension (σ), which is proportional to the contact length perimeter (σ ∝ L), becomes significant compared to the limited force that the robot can exert (F ∝ L^2^)^21^. Thus, small robots typically avoid large hydrophilic structures, which could provide more momentum, due to high surface tension impeding the leg rowing.

The other type of small-scale robots swings hydrophilic pads, which passively bends, back and forth only underwater to circumvent surface tension effects^22-26^. These robots lose momentum when recovering the pad underwater for the next stroke. Rhagovelia, in contrast, overcomes these limitations by using hydrophilic, reversibly actuating oars for effective drag-based propulsion, even at smaller scales than robots. Thus, designing artificial morphing fans inspired by Rhagovelia could address the limitations of small-scale robots, improving locomotion performance in turbulent waters where no small semi-aquatic robot has yet succeeded.

Here, we reveal that Rhagovelia possesses a fan with a unique flat-ribbon microarchitecture, enabling ultrafast capillary actuation. This design allows the barb to exhibit divergent rigidity in orthogonal directions, facilitating both elastocapillary morphing and effective force production during a propulsive stroke. This rapid, reliable self-spreading mechanism, independent of muscle control, enhances thrust production via unsteady vortices and capillary waves. In addition, the fan’s collapse helps to minimize surface tension impedance during the recovery stroke. We designed a robotic self-spreading fan modeled after Rhagovelia’s fan to explore these locomotion benefits. Our experiments show that a rowing robot with these fans performs significantly better than robots without fans, which agrees with finding in ripple bugs. These findings align with observations in insects, where intact fans provide superior maneuverability and endurance in unsteady waters compared to disabled fans. Thus, the ultrafast, passive actuation of Rhagovelia’s fan seems fundamental for effective thrust production, agile maneuvering, and sustained performance in turbulent aquatic environments.

## A self-morphing propulsor

SEM images reveal that Rhaghovelia’s fan comprises of numerous flat, ribbon-like barbules and barbs (Fig. 1a-c). The barbs interconnect and branch, while the barbules are uniformly distributed along both sides of each barb (Fig. 1c). Rhagovelia’s claw, smooth and flat, resembles a seam ripper tool (Fig. 1b). Both the fan and the claw attach to the tip of the leg. We observe that the fan of an isolated leg fully spreads underwater only when the claw is unfolded (Supplementary Fig. 1). Upon removal from the water, the fan collapses and adheres to the claw, forming a sharp tip, with no noticeable water dripping (Supplementary Video 1).

To demonstrate that Rhagovelia’s fan spreads due to capillary forces independent of muscle/claw, we isolated a fan from a specimen. Using a single human hair, we placed and removed the isolated fan from a water droplet (Fig. 1e). The fan fully spread in ∼10 ms upon contact with water, with barbules spreading even faster, at just 1 ms after water contact (Fig. 1e, Supplementary Video 1). In agreement we found that the theoretical spreading time for barbules, using the elastocapillary equation, has a similar order of magnitude to Rhagovelia’s fan (Supplementary Fig. 2, and Supplementary Table 1). The fan rapidly collapsed upon removal from the water, also within ∼10 ms. Experiments with a fixed leg and fan, gradually raising the water level, showed that the fan fully spreads only when completely submerged (Supplementary Fig. 3, Supplementary Video 1). Additionally, a wet fan in a droplet remains spread after drying (Supplementary Fig. 1). These experiments confirm that the isolated fan acts as an ultrafast capillary self-actuator without claw mediation, requiring full submersion or removal from water for complete spreading or collapse.

## Fan spreading during rowing and swimming

We analyze Rhagovelia’s rowing performance to understand fan spreading rates. Individuals initiate a leg stroke by reducing the gap between their hydrophobic leg and the water surface (Fig. 1f, Supplementary Video 1). This triggers the fan to open, reaching full spread in ∼10 ms and staying open for most of the stroke (∼50 ms, Fig. 3a-d). At the stroke’s end, they remove the fan from the water, closing it in ∼10 ms. The claw’s movements synchronize with the fan’s opening and closing. During the power stroke, the leg maintains contact at the tip throughout the inter-stroke period. These observations indicate that the fan spreading and closing dynamics during rowing mirror the passive behavior of an isolated fan.

**Fig. 2.**
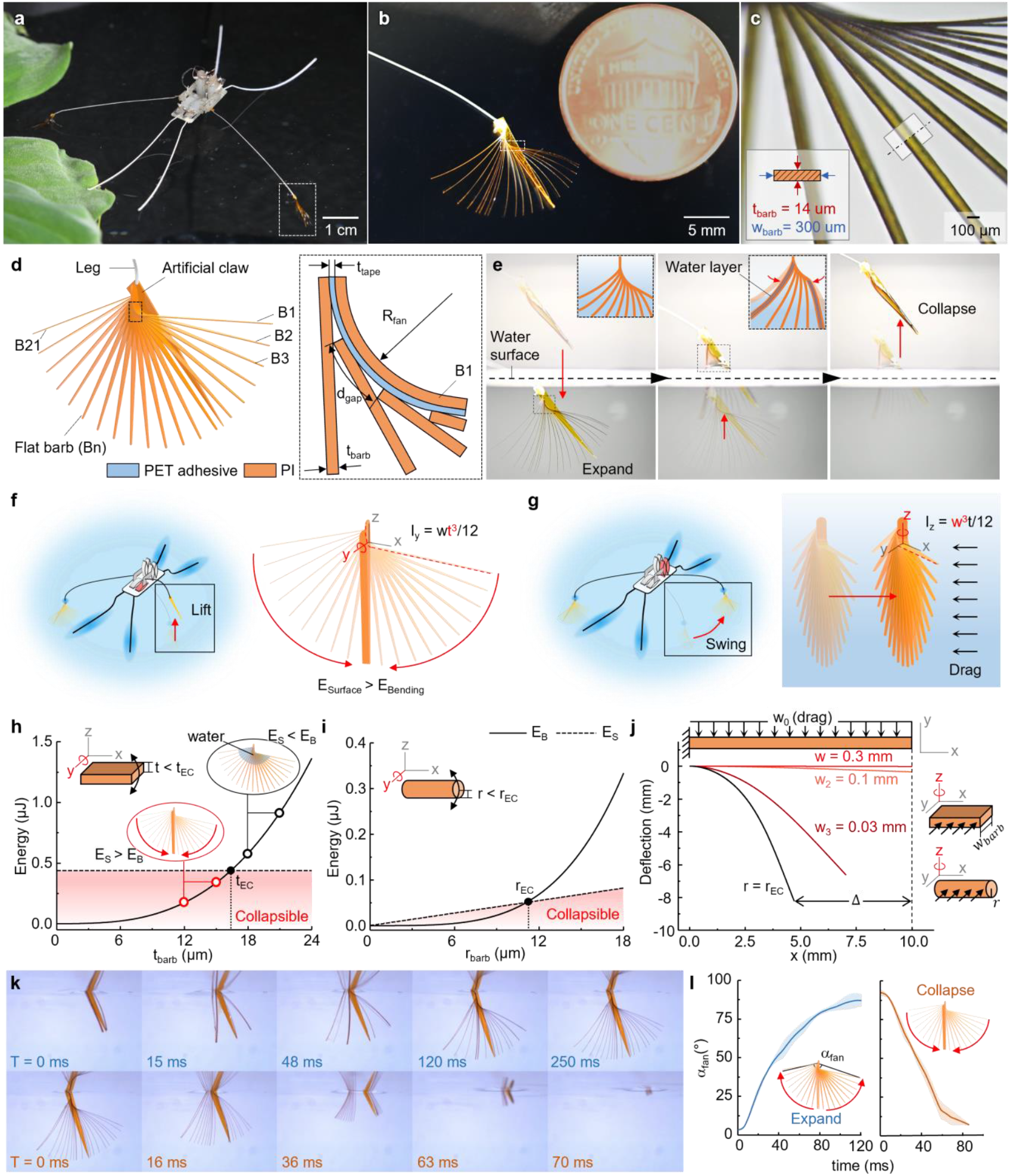
Design and mechanism of robotic fan driven by elastocapillarity. **a**, The 0.23-g semi-aquatic robot, Rhagobot, with self-morphing fans at the ends of its middle legs. **b**, The 1-mg robotic fan expanded underwater. **c**, Close-up view of the fan showing flat-ribbon barbs that mimic the structure of the biological counterpart. The thickness and width of the barbs are optimized to resist drag and achieve capillary morphing. **d**, Schematic of the fan consisting of 21 barbs (B1 ∼ B21) and an artificial claw. The radius of curvature (R_fan_) and distance between the start of each barb (d_gap_) determine the fan-like structure. The thickness of the barb (t_barb_) and PET adhesive (t_tape_) determine whether the structure will collapse. **e**, Sequence of images showing the fan’s expansion and collapse. The fan retains its expanded shape under water (left), starts collapsing due to capillary force upon elevation from the water (middle), and completely collapses in the direction of the artificial claw above the water (right). Schematic of the robot lifting its leg from the water **(f)** and swinging its leg underwater **(g)**, demonstrating the required low bending stiffness around the y-axis **(f)** for capillary-driven collapse and high bending stiffness around the z-axis **(g)** to prevent bending due to drag. Bending (solid line) and surface (dashed line) energy as a function of thickness and radius (t_barb_, r_barb_) of barb with rectangular cross-section **(h)** and circular cross-section **(i)**. The thickness must be less than the elastocapillary thickness and radius (t_EC_, r_EC_) to enable capillary force-driven collapse. See Supplementary Table 2 for details of calculation. **j**, Comparison of deformation in capillary-driven beam with circular and rectangular cross-sections under drag forces. **k**, Sequence of images showing the robotic fan expanding when entering the water (top) and collapsing when exiting the water (bottom). **l**, Angle (α_fan_) profile of a fan during expansion and collapse as it enters and exits the water.

**Figure 3.**
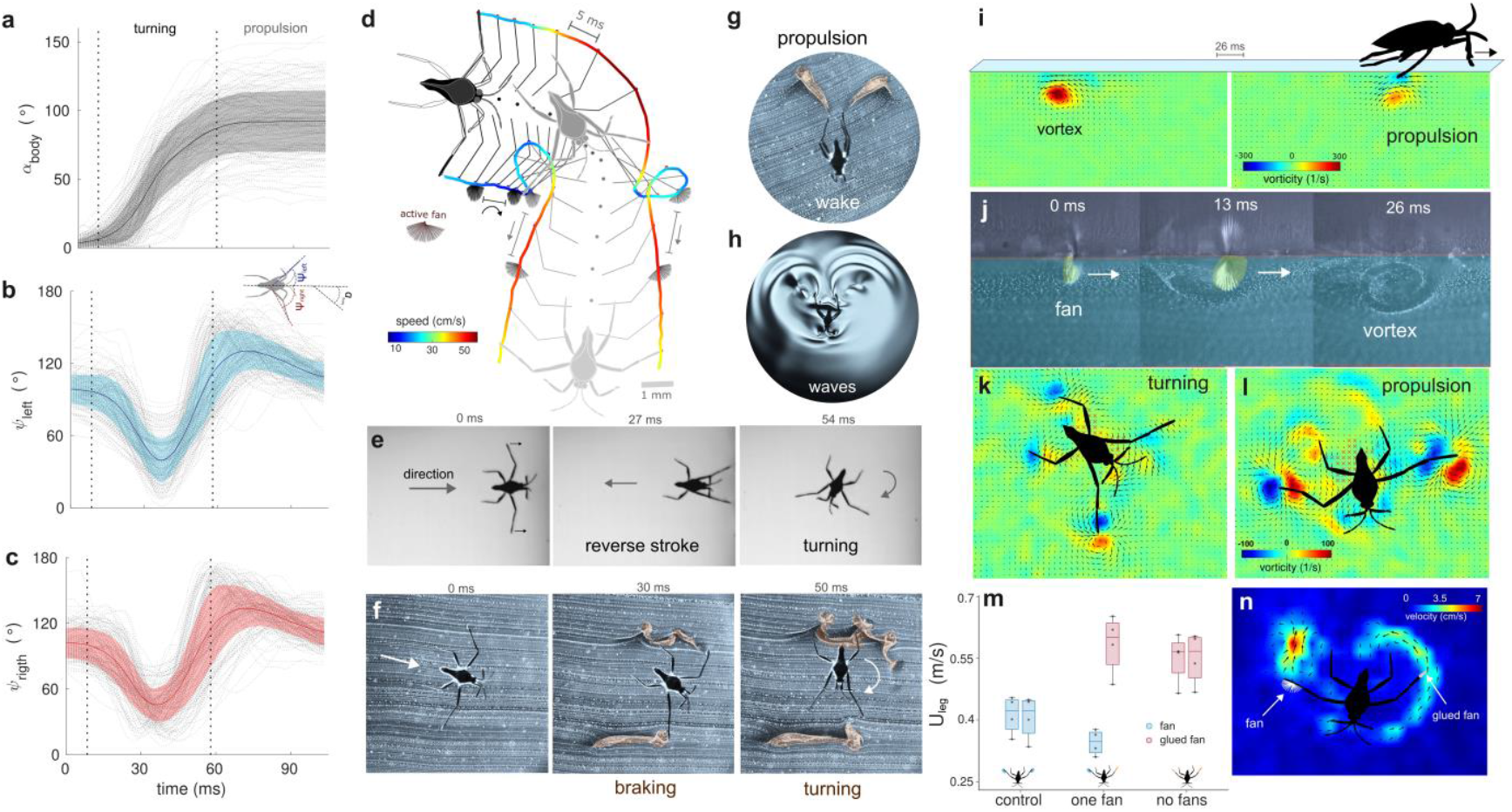
Rhagovelia’s kinematics and hydrodynamics. Time series of the body angle **(a)**, left leg **(b)**, and right leg **(c)** during turning maneuvering and propulsion. Dashed lines represent each trial (N=159). The colored line and shaded region represent the mean and one standard deviation. **d**, Detailed illustration of turning and propulsion of an individual showing when the fan is spread. The timelapse between points is 5 ms, and the color bar represents speed. **e**, Braking maneuver using a symmetric reverse stroke and turning. **f**, Vortical wake produced during braking and turning. Vortical wake **g**, and capillary waves **h**, produced during propulsion. Side view of vortex produced by Rhagovelia during propulsion showing the vorticity field **i**, and flow visualization **(j)**. Notice the spread fan orientation in J. Top view of turning **(k)** and propulsion **(l)** showing the vorticity field. **m**, Leg speed during propulsion for individuals (N=4) with both fans intact (control), one fan inactivated (one fan), and both fans inactivated (no fans). Error bar represents mean ± one s.d. **n**, Velocity field using PIV of an individual with one fan inactivated. Notice that the intact fan induces higher flow speed than the inactivated fan.

We assess the claw’s role during Rhagovelia’s underwater swimming. Folding a submerged self-spread fan with the claw is more challenging than opening, as the claw must actively force the fan to fold. Consequently, the fan remains partially spread rather than fully closed during swimming (Fig. 1j-i, Supplementary Video 1). These findings reveal that Rhagovelia exerts both active and passive control over fan dynamics. Fan closing relies on passive elastocapillary forces when removed from the water or active via claw-muscle action when fully submerged. Fan opening is predominantly passive, but the claw can act as a gate, controlling the release of the fan underwater.

## Robotic fan design and passive operation

We develop an artificial fan to investigate the mechanical advantages of a flat-ribbon structure over a cylinder hair-like structure, focusing on elastocapillary self-morphing and drag-force production. This fan, attached to a bio-inspired insect robot called “Rhagobot,” is placed at the tip of each rowing leg. Measuring 10 × 5 mm^2^ and weighing ∼1 mg, the fan folds and spreads passively via elastocapillarity (Fig. 2a). It consists of 21 flat ribbon shaped barbs with a claw-like structure (5 mm in length), and its size is ∼20 times larger than Rhagovelia’ s fan (Fig. 2b). Each barb is made from an optimal 16 μm thickness of Polyimide (PI) film, processed by a UV laser (Fig. 2c and Supplementary Fig. 4). The first barb (B1) is connected with 1 μm Polyethylene terephthalate (PET) tape to each other barb, facilitating equidistant spreading (Fig. 2d).

The artificial claw at the fan’s base enables the barbs to collapse under surface tension in the desired direction. Rapid closing and spreading occur when the fan is removed from or introduced to water, respectively (Fig. 2e). This capillary-driven passive operation allows the fan to function during the Rhagobot’s stroke cycle without needing an actuator. This elastocapillary morphing offers two advantages: first, the fan’s collapsibility reduces surface area, minimizing surface tension during the recovery stroke (Fig. 2f), and second, the fan’s spreading facilitates drag-based propulsion during the power stroke retaining its shape during the power stroke (Fig. 2g).

## Optimizing elastocapillary and drag forces

We design artificial barbs with a flat ribbon shape to optimize the fan for elastocapillary collapse and drag resistance by ensuring different stiffness levels in two orthogonal directions. For effective elastocapillary folding, the barb must bend around the y-axis due to surface tension (Fig. 2f). The surface energy needs to exceed the bending energy, which depends on the moment of inertia around the y-axis, a function of *t*^*3*^ (where *t* is the barb’s thickness). By adjusting the barb thickness, we reduce the bending energy below the surface energy, facilitating elastocapillary folding. To maximize drag-based propulsion, the barb deformation must be minimized during the power stroke. Drag-induced deformation centers around the z-axis, with the moment of inertia around the z-axis being a function of *w*^*3*^ (where *w* is the barb’s width, Fig. 2g). By optimizing both thickness (*t)* and width (*w)* through modeling, the flat ribbon shape achieves both elastocapillary-driven morphing and maximum drag force.

We quantify the mechanical advantages of the flat-ribbon shape over a cylindrical-shaped barb. By comparing the bending and surface energies of the barbs at the ends (B1 and B21) when unfolded and folded, we determine the maximum thickness for elastocapillary bending to be *t*_*EC*_ = 16.3 μm (Fig. 2h, Supplementary Fig. 5 and Supplementary Table 2). For a cylindrical barb, the maximum radius for collapse is *r*_*EC*_ = 11.1 um (Fig. 2i). While the thickness of the flat ribbon shape remains fixed at *t*_*EC*_, adjusting the width (*w*_*fan*_) minimizes deformation due to drag while maintaining collapsibility. Increasing *w*_*fan*_ up to 0.3 mm results in negligible deformation (*Δ*_*barb*_), maximizing drag utilization (Fig. 2j). In contrast, a cylindrical barb with *r* = 11.1 μm undergoes 70% deformation (*Δ*_*barb*_), reducing the drag-receiving surface area and thus, propulsion force. Increasing *r*_*EC*_ to minimize deformation increases the moment of inertia around both the z-axis and y-axis, compromising collapsibility. The energy balance indicates that not only the artificial fan but also the Rhagovelia fan is collapsible, as the surface energy is three orders of magnitude higher than the bending energy of the biological fan (Supplementary Table 3 and 4, and Supplementary Fig. 6).

The optimized flat ribbon design ensures the artificial fan is both collapsible and maximizes thrust production. Our design suggests that the flat-ribbon microstructure of Rhagovelia’s fan enhances rowing on the water surface. Observations of the artificial fan, lifted and lowered by the Rhagobot, show that it opens in ∼120 ms and collapses in ∼80 ms in water (Fig. 2k and 2l, and Supplementary Video 2). This finding agrees with the theoretical spreading time ∼O (300 ms) for artificial fan calculated using the elastocapillary equation (Supplementary Fig. 2g, and Supplementary Table 1 and 5).

## Controlled maneuvers and vortex hydrodynamics

Rhagovelia exhibits an average turning angle of 90°, a turning rate of 4.2×10^3^°/s, and a turning duration of 50 ms, similar to a fly (see Discussion and Fig. 3a-f). A turn begins when an individual moves both legs forward in a reverse stroke, with only one leg and its fan interacting with the water. This asymmetric braking turns the body towards the dragging leg. During the subsequent power stroke, both legs and fans engage synchronously, producing symmetric vortical wake and capillary waves (Fig. 2d and 2h). Rhagovelia achieves higher turning angles with fast braking control. In a reverse stroke using both legs and fans, this action induces a complex wake of interconnected dipolar vortices resembling a Kármán vortex street (Fig. 2g). This sudden change in direction helps avoid obstacles in fast, unsteady streams, evade predators, or escape unwanted mates. We calculate that adult Rhagovelia, with a mass of 2.1±0.4 mg, traveling with a speed and acceleration of ∼0.2 m/s and ∼12 m/s^2^, respectively, exerts a force of 26 µN and a momentum of 4.5×10-7 kg m/s during propulsion.

When turning, Rhagovelia induces a wake of dipolar vortices with the fan in contact with the water (Fig. 3a). These vortices have vorticities up to ∼100 s^-1^ and flow velocities up to ∼20 cm/s. During propulsion, vortices show similar vorticity, but the flow velocity doubles (Fig. 3a, Supplementary Fig. 7). Side-view PIV reveals that a propulsive Rhagovelia fan generates a strong vortex with a vorticity of ∼300 s^-1^ and a flow velocity of ∼70 cm/s. Tip vortices are shed and remain connected to the fan during a propulsion stroke. Capillary waves form during propulsion, along with bow waves at the front of the body (Fig 3h and Supplementary Video 1). Constructive interference occurs when waves from each leg converge at the back (Fig. 3h, Supplementary Video 1).

## Active and inactive fans

To understand the role of fans in Rhagovelia during the propulsive stroke, we conduct a controlled experiment. We film four adults with intact fans rowing on the water surface using a high-speed camera. Next, we inactivate one fan on each individual with Loctite super glue and dust the leg tip with lycopodium particles to maintain hydrophobicity (Supplementary Fig. 8). We then film these individuals rowing with one inactivated fan and one intact fan. Finally, we inactivate the remaining fan on each individual and film them rowing again.

Legs with intact fans consistently show lower tip speeds than legs with inactivated fans, regardless of the treatment stage (control, one fan inactivated, two fans inactivated, Fig. 3m and Supplementary Table 6). This demonstrates that intact fans significantly enhance momentum transfer to the water via drag. PIV analysis supports these findings, showing that legs with intact fans during a propulsive stroke double the maximal flow speed compared to legs with inactivated fans.

## Rhagobot’s actuation and control

For controlled leg movement in small-scale robotics, the Rhagobot uses four independent, lightweight shape memory alloy (SMA) actuators (17 mg) as shown in Fig. 4a. Actuators R1 and L1 manage the swing input, while R2 and L2 control the lift input, enabling robust horizontal and vertical leg movements (Fig. 4b). Each leg passes through holes in the swing and lift inputs, allowing it to swing horizontally or lift vertically (Supplementary Fig. 9a). A film guide attached to the leg and frame prevents unwanted leg rolling. Attaching an artificial fan to a leg tip demonstrates that the actuators can swing the open fan underwater without the fan bending and lift it from the water, achieving passive elastocapillary collapse (Fig. 4b). Design parameters specify the leg rotation angle (*θ*_*leg*_), with a stroke (δ) of the actuator yielding 80° for swinging and 40° for lifting (Supplementary Fig. 9a). By sequencing the actuators, the Rhagobot can perform various leg motions such as the power stroke, recovery stroke, reverse stroke, liftoff, and touchdown. These motions enable forward motion, braking, rotation while advancing, and rotation in place with the different signal inputs (Fig. 4c and Supplementary Fig. 9b).

**Fig. 4.**
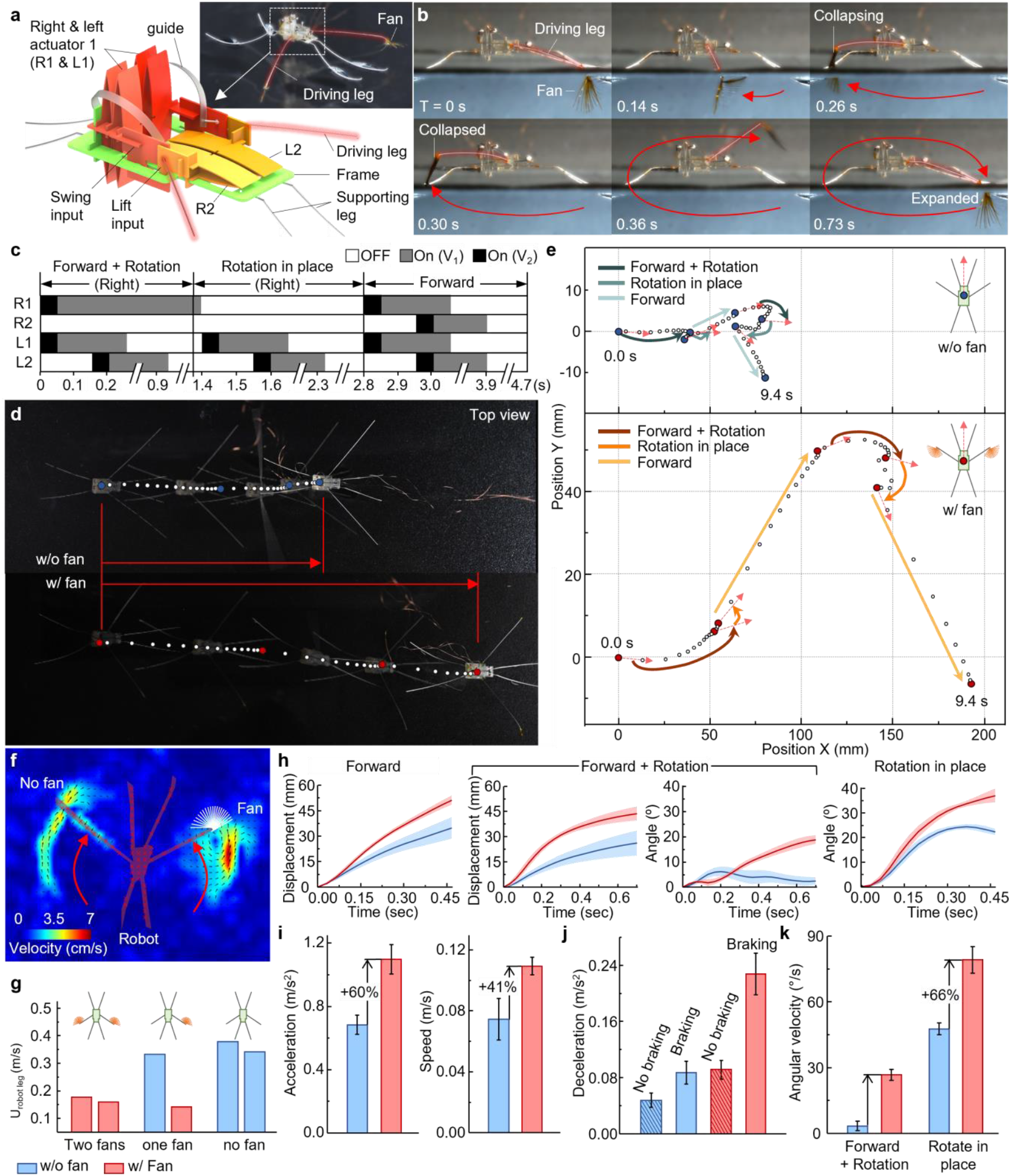
Rhagobot control, performance, and agility. **a**, CAD model of a rhagobot driven by four SMA-based actuators. **b**, Sequence of images showing the right side of the robot performing a rowing motion with a self-morphing artificial fan. **c**, Driving voltage sequence of the four actuators (R1, R2, L1, L2) to achieve three distinct motions: rotation while advancing, rotation in place, and forward motion. A high voltage (V_2_) induces full actuator stroke within 50 ms, and a lower voltage (V_1_) maintains the actuator’s extended state without overheating the SMA. **d**, Comparison of three sequential forward motions on the water surface with and without a fan. **e**, Trajectory of the robot with and without fan performing three motions: rotation while advancing, rotation in place, and forward motion on the right and left sides. **f**, Velocity field using PIV to compare a swinging leg with a fan and a superhydrophobic coated leg. **g**, Leg speed comparison for three robot configurations: both legs equipped with fans, only one leg with a fan, and neither leg with a fan. **h**, Distance and body angle profiles of robots with and without fans. **i-k**, Comparative analysis of acceleration, speed **(i)**, deceleration **(j)**, and angular velocity **(k)**, demonstrating the superior performance of the robot equipped with fans.

## Rhagobot performance with and without fan

To assess the impact of the artificial fan on locomotion, we construct two types of robots. The first robot has driving legs coated with a superhydrophobic material for surface tension-based propulsion, similar to Geriidae water striders. The second robot has our hydrophilic artificial fan on the driving legs for drag-based propulsion in water, mimicking Rhagovelia (Fig 2a and 2b). We apply identical input voltage signals to both robots, resulting in forward and turning motions. Robots with fans travel farther and make sharper turns compared to those without fans (Fig. 4d and e, Supplementary Video 2). Despite receiving the same input energy, the fan-equipped robot gains more momentum in water. PIV analysis shows the fan increases maximum flow speed by 33% and vorticity by 24% during propulsion (Fig. 4f, Supplementary Fig. 10).

Leg speed during a stroke confirms that the fan generates stronger momentum. Comparing three configurations – both legs with fans, one leg with a fan, and neither leg with a fan – the legs without fans consistently show high speed (indicating less resistance in water and consequently less thrust, Fig. 4g). This aligns with experiments on Rhagovelia with and without fans (Fig. 3m-n, Supplementary Video 1). Fan-equipped robots achieve greater momentum, enhancing distance traveled and turning angle (Fig. 4h). The fan increases the forward acceleration and speed of the robot by 60% and 41%, respectively (Fig. 4i). The average and maximum speed of the robot with fan are 12.14 cm/s (1.47 BL/s) and 16.16 cm/s (1.96 BL/s), respectively.

## Fan-assisted robot maneuverability

The fan-induced thrust increases forward speed and allows rapid braking as well (Supplementary Video 2). Without the fan, lowering both legs during forward motion does not sharply reduce speed due to low shear force on the superhydrophobic legs (Fig 4j). Lowering one leg into the water while moving forward enables yaw turns centered on the fan. For faster rotation, swinging both legs in opposite directions induces rapid on-the-spot yaw rotation. The robot’s average and maximum rotation speed for on-the-spot rotation are 87.1 °/sec and 206 °/sec, respectively, with the fan (Fig 4k). The Rhagobot exhibits agile turning and faster speed among small-scale semi-aquatic robots. Rowing with large hydrophilic pad has traditionally been feasible only in large-scale robots, which possess sufficient power output to overcome surface tension (Fig. 5a and 5b). In contrast, Small-scale robots typically rely on surface tension-based propulsion, which is limited until the water surface is broken. The other propulsion method of small-scale robots involves the back-and-forth movement of passive flaps underwater, which leads to a reduction in a speed during recovery phase. The self-morphing elastocapillary fan enables the Rhagobot to generate greater momentum by maintaining its large surface area underwater and avoiding speed reduction in recovery phase by retracting the collapsible fan from the water (Fig. 5a and Supplementary Table 7).

**Fig. 5.**
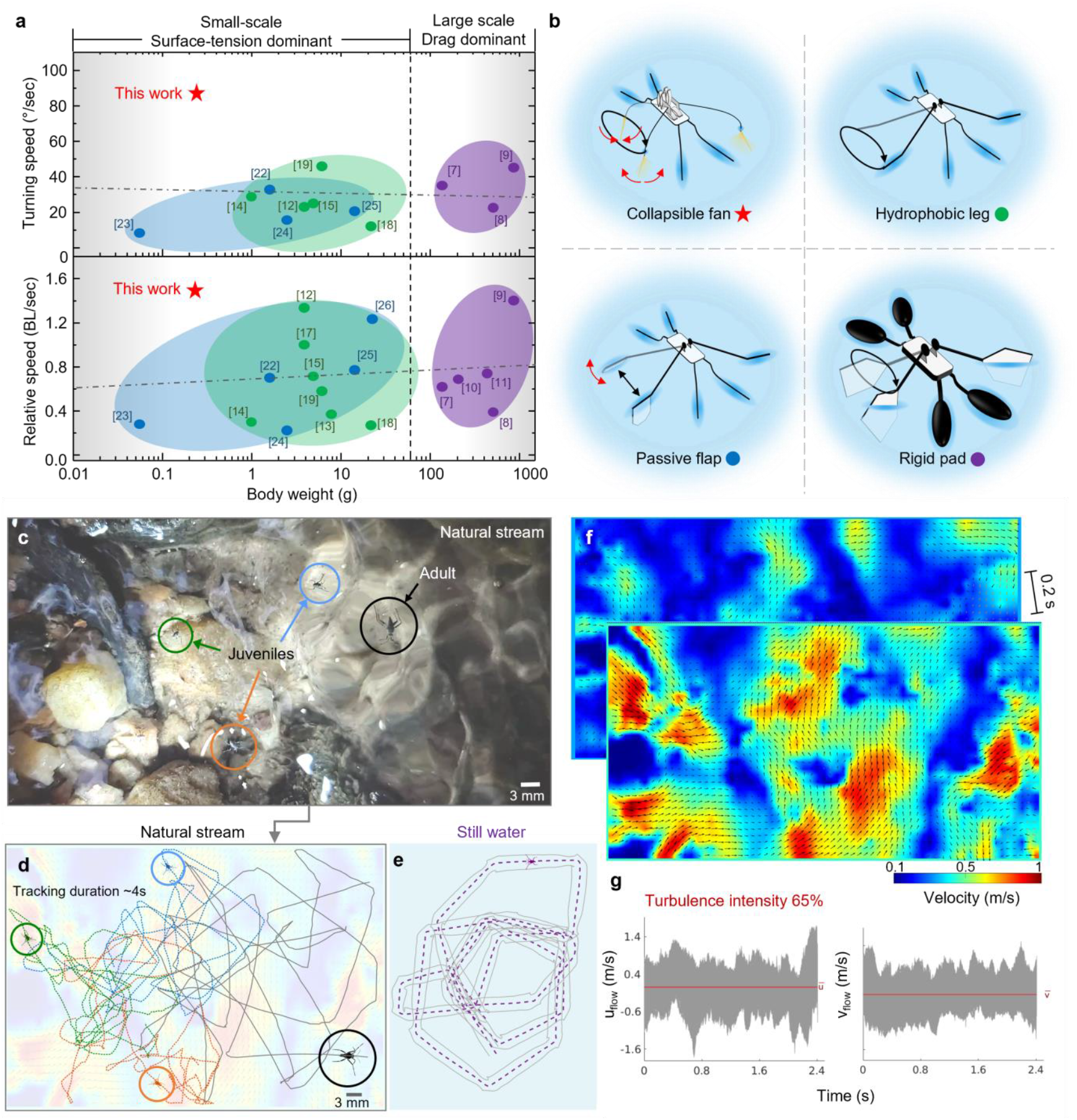
Semi-aquatic robot comparison and flow conditions in natural streams. **a**, Turning speed and relative speeds comparison of semi-aquatic robots relative to body weight. Dashed line represents a linear fit of the robot performance data. **b**, Schematics showing four different propulsion mechanisms of semi-aquatic robots: rowing with a collapsible fan (top left), rowing with a thin hydrophobic leg (top right), swinging a passive flap back and forth (bottom left), and rowing with a large rigid pad (bottom right). See Supplementary Table 7 for details of the robots. **c**, Video frame showing one adult and three juveniles moving on a natural stream. Colors represent each individual tracking. **d**, Irregular trajectories followed by those individuals during ∼4 seconds. **e**, Regular path followed by an individual rowing in still water. **f**, Velocity fields of flow condition that those individuals experience *in situ*. Notice the dramatic change in flow speeds after 0.2 s. **g**, Time series of U_flow_ and V_flow_. Turbulent intensity (ratio between flow speed variation and average flow speed) was ∼65%. See text for details.

## Turbulence in natural streams

To evaluate the natural flow condition that ripple bugs experience in the wild, we perform PIV analysis of the surface of streams where these insects are found. Lycopodium particles were used to seed the flow. We found that ripple bugs withstand streams with flow speeds up to 1.6 m/s and turbulence levels (i.e. the ratio between the root-mean-square of velocity fluctuations and the mean velocity) of 65% (Fig. 5d-g). Juveniles and adults can be found sharing similar flow perturbations (Fig. 5c-d). Trajectories followed by Rhagovelia in natural streams are irregular, which contrast with the regular ones followed by individuals in still water (Fig. 5e). This high levels of turbulence indicate that fast maneuvering and propulsion, mediated by a rapid fan actuation is fundamental for Rhagovelia to stand such adverse flow conditions.

## Rhagovelia’s non-stop rowing activity

In order to understand Rhagovelia’s rowing activity we monitor dozens of individuals, from first instar to adults, using a GoPro Hero camera. We found that these individuals are active night and day, rowing non-stop during most of their life span. Some individuals survived up to ∼8 months (Supplementary Video 1). We observe that these insects only rest during molting, but only for a few minutes or occasionally while feeding and cleaning. Males stop rowing during mating too, because they ride on the female’s back, but females keep rowing all the time despite the extra weight. This outstanding endurance shown by Rhagovelia agrees with the increased rowing performance added by their elastocapillary fans.

## Discussion

### Adaptations for unsteady aquatic niches

Natural streams are characterized by unsteady dynamics and abundant prey^27^. These habitats pose significant challenges for terrestrial insects, being difficult, dangerous, and energetically costly to transit. In-situ field PIV analysis reveals that Rhagovelia moves in waters with flow speeds up to 1.6 m/s and turbulence levels of 65% (Fig. 5f and 5g). Long-term lab timelapse observations uncover that Rhagovelia rows continuously, day and night, stopping only briefly for feeding, cleaning, or molting (Supplementary Video 1). At an average speed of 0.14 m/s, Rhagovelia can travel per day∼13 km, which is similar than the long-distance flights exhibited by fruit flies^28^. Impressively, Rhagovelia bugs can live in the lab up to eight months (Supplementary Video 1), thus covering a total distance of ∼3000 km Rhagovelia’s unique fans enable ultrafast passive actuation, facilitating agile turning and braking control, and effective propulsion via vortex generation, essential for continuous foraging in highly perturbed flows.

### Fan structure and function

The anisotropic barbs and barbules of Rhagovelia’s fan promote rapid passive spreading and collapse, akin to a paintbrush interacting with liquid. Synthetic fan experiments show that this flat-ribbon geometry provides divergent stiffness in orthogonal directions, balancing elastocapillary spreading and drag-based thrust. The fan’s minimal water retention aligns with observations in paintbrushes with flexible fibers and dandelion seeds ^29^. Rhagovelia keeps the tip of its leg on the water to avoid the extra load of a wet fan (Supplementary Fig. 3), contrasting with Gerridae water striders, which elevate their hydrophobic legs during recovery strokes^30^. The barbs’ slightly depressed middle region forms channels that may facilitate water conduction (Supplementary Fig. 11), resembling structures in desert plants and invertebrates for water harvesting^31^.

### Active and passive control mechanisms

Rhagovelia exhibits both passive (elastocapillarity) and active (claw-muscle) control over fan movement. The duration of fan spreading and collapse is similar in both cases. However, muscle-driven fan closing during swimming often leaves the fan partially spread, increasing drag during recovery strokes. Nonetheless, active control mechanisms allow precise locomotive control during unexpected challenges. Other water bugs, such as *Ptilomera sp*^*30*^ and *hallobates spp*^*32*^, have specialized hair brushes on their middle legs, which may also self-spread underwater through elastocapillary action. Comparative studies of these structures could provide insights into the evolutionary trends and locomotive functions of hair-like structures in water striders.

### Turning like a fly

Rhagovelia’s locomotion on the water surface mirrors the acrobatics of flying fruit flies during banking maneuvers. Rhagovelia can turn 90° at a rate of 4200 °/s, in 50 ms, metrics that are similar to the fruit fly’s quick saccades of 93° turns at rate of 5000°/s in 49 ms^33,34^. While flies rely on continuous wing flapping for sharp yaw rotations^35^, Rhagovelia achieves these maneuvers with a single reverse stroke. Compared to semi-aquatic whirligig beetles, Rhagovelia’s turning rates are 50% faster, and its stroke frequency (∼20Hz) is 2.5 times lower^36,37^. Whirligig beetles are limited to curved circular trajectories in slow, low-turbulence waters, while Rhagovelia performs sharp turns at high rates with low stroke frequencies in unsteady flow conditions. Moreover, whirligig beetles, in contrast with ripple bugs, are seen resting for prolonged periods of time during the day and night (VMOJ, personal observation).

### Vortices and waves

Aquatic paddling birds generate thrust via lift using their feet as inverted delta wings^38^, suggesting a similar lift-based thrust mechanism in Rhagovelia fan-based propulsion. Moreover, Rhagovelia’s wake during rowing comprises multiple vortices resembling tip vortices on flying wings during propulsion (Fig. 3g and 3f, and Supplementary Video 1), warranting further hydrodynamic investigations. Rhagovelia also produces capillary waves during leg strokes, similar to Gerridae species and whirligig beetles^20,39^. These capillary waves might serve locomotion, communication, or sensing functions, both in individuals and collectives, presenting intriguing avenues for ongoing research.

### Fan-assisted robotic maneuvers

Simple and lightweight passive actuation structures are essential for microrobots, eliminating the need for active actuators. Strategies such as passive wings^40^, flaps^22,26^, and alignments^41^ have been widely used to interact with the environment. This concept, known as embodied intelligence^42^, is exemplified by Rhagovelia’s fan. Inspired by this fan, we engineered an ultralightweight, 21-layer robotic fan (∼1 mg) using micro multilayered fabrication. This collapsible and rigid bio-inspired fan balances structural elasticity, capillary force, and drag, enhancing interfacial microrobot performance. Our robotic fan, with its collapsibility and rigidity, effectively uses drag to maximize momentum while minimizing surface tension interference, resulting in faster and more efficient movements compared to other small-scale robots.

The Rhagobot employs four independent lightweight actuators (∼17 mg), allowing each leg to be controlled with two degrees of freedom (DOFs). These actuators enable the Rhagobot to perform various leg motions, such as power stroke, recovery stroke, reverse stroke, liftoff, and touchdown, resulting in controlled rapid maneuverability with a maximum forward and turning speed of approximately 1.96 BL/s and 206 °/sec, respectively. Although our current design is tethered, future versions could be autonomous and capable of navigating rough, turbulent stream surfaces with self-actuating oars.

## Acknowledgments

VMOJ thanks Chris Sanford and KSU for supporting the beginnings of this project. S.B. acknowledges funding support from NIH MIRA Grant R35GM142588; NSF Grants PHY-2310691; CMMI-2218382; CAREER iOS-1941933; and the Open Philanthropy Project. J.-s. K. acknowldeges funding support from National Research Foundation of Korea grants funded by the Korea government (grant nos. RS-2021-NR061649 and RS-2024-00411660).

## Authors contributions

V.M.O.-J. conceptualized the research and designed the animals’ experimental framework. D.K. and J.-S.K. contributed to the design of the artificial fan and semi-aquatic robot. D.K., C.K., and J.-S.K. developed the robot’s experimental design. V.M.O.-J., D.K., J.-S.K., and S.B. contributed to the research methodology. D.K., C.K., and J.-S.K. conceived the theoretical model of the robotic fan. S.K. and S.B. analyzed the fan’s physical properties. V.M.O.-J conducted fieldwork and fluid dynamics analysis. S.B. and J.-S.K. provided funding for the project and supervised the research. V.M.O.-J., D.K., J.-S.K., S.K., and S.B. contributed to writing, reviewing, and editing of the manuscript.

## Materials and Methods

### Passive actuation

We remove fans from two Rhagovelia adults using dissecting tools and an optical microscope. Individuals were previously placed in 70% ethanol alcohol. Removed fans were washed with water to remove all alcohol content. Then, we use human hair to manually take an individual fan from water to the air and gently place the fan on a droplet of water attached to a microscopic slide. We filmed from above the spreading (n=9) and closing (n=9) of the fan when it was manually placed in and out of water using the hair. A high speed camera (Nova S6, Photron, Inc.) at 2000 frames/s with a 4x microscope objective was used. We manually digitized the outer tips of the fan and the root from those video sequences using DLTdv8 for Matlab^43^. We used those digitized points to calculate the spreading angle over time.

### Fan and claw spreading during rowing

We filmed sideview three Rhagovelia adults while rowing at the interface using a high speed camera at 2000 frames/s. To have a clear image of the fan we tilted the plexiglass container in a way that the water meniscus was minimal. We filmed 13 trials from all individuals while spreading and closing their fans. From those video sequences the outer tips of the fan, as well as the tip and root of the claw were digitized. We used those digitized points to calculate the fan and claw spreading angles over time.

### Fan and claw spreading during swimming

Two Rhagovelia adults were filmed at 2000 frames/s swimming to understand the active actuation of the fan underwater. We filmed 5 and 6 trials closing and spreading the fan via the claw, respectively. Fan and claw spreading angles over time were calculated as they were explained before.

### Turning performance

We label manually 277 frames from varied of the sampled videos (Supplementary Fig. 12). From those digitized points we calculated the body angle, each leg stroke amplitude and the head speed over time.159 turning trials of 57 individuals (adults and juveniles) were filmed at 749 (Fastec HiSpec 4), 1000 (Mini Ax200) and 4352 (Chronos 2.1) frames/s. The tips of the head and abdomen, as well as the tip and root of both legs were digitized using DeepLabCut (http://www.mackenziemathislab.org/deeplabcut). We label manually 277 frames from varied of the sampled videos. From those digitized points we calculated the body angle, each leg stroke amplitude and the head speed over time.

### Particle image velocimetry

A laser (Opto Engine LLC, 532 nm, 5 W) was used illuminate plastic beads (50 µm) at the surface of the water. We filmed at 1000 frames s^−1^ (Fastec HiSpec 4). Velocity and vorticity fields were calculated using PIVlab (http://william.thielicke.org/PIVlab). An interrogation window from 64 to 32 pixels^2^, excluding those vectors with standard deviation greater than 5, was used.

### Wake Visualization

We filmed (1000 frames/s) the wake of 3 individuals while moving on a flume (∼5 cm/s) over a sheet of hydrogen bubbles at the interface. Hydrogen bubbles were generated using electrolysis, a classic flow visualization method.

### Capillary Waves

We filmed (1000 frames/s) the capillary waves of two Rhagovelia individuals while rowing on water. In order to see the waves we use a pinspot light source placed at 45° with the horizontal and pointing down.

### Rhagovelia in natural streams

Activity of Rhaghovelia water striders was filmed at 120 fps (S10 Samsung Galaxy) in natural streams in Kennesaw, Ga, close to Kennesaw State University. Centroid of individuals was tracked manually using DLTdv8. Lycopodium particles were added to the stream and filmed from above. We use those videos to extract the velocity fields as mentioned before.

### Day and night activity

A GoPro Hero 11 was used to film (time lapse each 10 s) the activity of ∼20 Rhagovelia water striders in the lab from hatching (first instar) to adult (four instar). A Tetra Whisper Power Filter 10 Gallons was used to keep the water clean and running in the plastic container. Each two days individuals were fed with fruit flies or springtails. A LED lamp (VILTROX L116T, set up at 7% intensity) was used to provide dim light during night time.

### Intact and disable fan individuals

We inactivate one fan on each individual with Loctite super glue and dust the leg tip with lycopodium particles to maintain hydrophobicity (Supplementary Fig. 8). Four adults were used for this experiment. Each individual was allowed to move on a paper filter to remove any water remaining on legs. Then each sampled individual was anesthetized by placing it momentarily on ice. A pipet tip with a drop of Loctite Super Glue was introduced at the tip of the leg of each individual to glued the fan. Then immediately the glued leg was dusted with lycopodium particles. We allow each individual to recover for 5 minutes. We repeated the procedure to glue and dust with lycopodium particles on the other leg. Three treatments resulted from this procedure: Individual rowing with both intact fans, with only one disabled fan, and with both disabled fans. We filmed from above each individual per treatment using a high speed camera at 1000 frames/s. We used DeepLabCut to digitize the head and the tip of the abdomen, as well as the tip and root of both legs. Tip speed of each leg was calculated using digitized leg tip points and following^44^. We use a two-way repeated measures ANOVA to compare the effect of fan (intact vs disabled) and leg treated (left or right) in the maximal speed reached by the leg tip during the power stroke. ANOVA test was performed in R V4.4.1 (https://www.r-project.org/)

### Artificial fan fabrication

The construction of the artificial fan involves an assembly of thin film layers, comprising twenty-one layers of 16 µm thick polyimide (PI) film and a single layer of 1 µm thick polyethylene terephthalate (PET) adhesive. The assembly process begins by aligning twenty PI film layers (B2 to B21) with a defined gap (d_gap_) of 150 µm between each (Supplementary Fig. 4a). This spacing allows for the subsequent attachment of the PET adhesive. The final layer, a PI film with an adhered PET layer (B1), is then aligned and placed on top of the exposed spacing of the layers (B2 to B21), (Supplementary Fig. 4b). Following the alignment, a rubber pad is placed over the layers, and uniform pressure is applied using a heat press (Supplementary Fig. 4c). The conditions set for this process are 10 MPa at 40°C for 30 seconds, effectively bonding the B1 layer to the underlying twenty layers (B2 to B21) through the PET adhesive.

After pressing, the entire layered structure is cut into a fan shape using a UV laser machine (spec.). The dimensions for cutting are set to a length of 1 cm and a width of 300 µm (Supplementary Fig. 4d). To deploy the compact structure into a fan shape, the B1 layer is partially wrapped around a 1.1 mm diameter rod, securing approximately half a turn (Supplementary Fig. 4e). This configuration allows the barbs (B2 to B13) to spread out tangentially, forming a semi-fan shape. To permanently set the bent shape of B1, the deployed assembly is heated in a 300°C chamber for approximately 10 minutes. Once the fan is removed from the fixed rod, B1 exhibits an increased radius of curvature compared to its initial configuration. The remaining barbs (B14 to B21) can be folded out at regular intervals to complete the fan’s fan-shaped configuration.

### Model of energy balance and large beam deflection for a robotic fan

To ensure that an artificial fan collapses via capillary force, the geometry of the fan must be determined by considering both surface energy and bending energy, specifically to select a maximum thickness of the barb. Although the fan has 21 barbs, we will consider only the energy of the two most bent barbs at the extremes, B1 and B21 (Supplementary Fig. 5a). We assume that if only the two extreme barbs are collapsible, then the added barbs in between will also collapse. If the surface energy (Es) of the two extreme barbs exceeds their bending energy (E_b_=E_b1_+E_b2_), then the two barbs will collapse into one. The definitions of bending energy and surface energy are as follows:

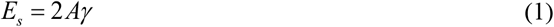

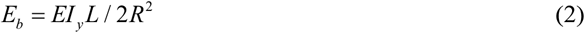

where γ is the surface tension of the liquid, and A is the surface area of one side of the barb. I is the second moment of inertia and R is the radius of curvature of the curved barb (I_lamella_=wt^3^/12, I_rod_=r^4^/4π). As the number of barbs between the two extremes increases, both bending energy and surface energy increase. However, the increase in surface energy is greater than that of the bending energy, because the increase in surface energy is a multiple of 2 *Aγ*, while the increase in bending energy is less than the initial E_b1_+E_b2_. Thus, by considering only the most extreme case of the two barbs, we can determine the minimum design criteria that allows a fan with 21 barbs to collapse.

To determine the maximum thickness of the lamella-shaped barbs that ensures collapsibility, we select thickness as a variable and find the maximum thickness that satisfies Es > Eb. This energy balance equation is given from (1) and (2) by the following cubic equation in terms of thickness (t):

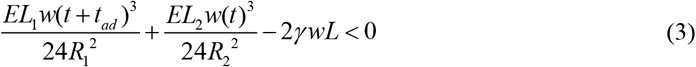

where L_1_ and L_2_ are the lengths of the curved barbs (B1 and B21), R1 and R2 are the radii of curvature of B1 and B21, and t_ad_ is the thickness of the adhesive (Supplementary Fig. 5b and 5c). The values of the parameters are reported in Supplementary Table 2. Thus, thickness (t) must be designed to be less than approximately 16.3 µm for the barbs to coalesce due to capillary force. A fabricated 12 and 14 μm thick barb, smaller than t_EC_, demonstrates collapsibility due to lower bending energy compared to surface energy. In contrast, experiments with two barbs of 18 and 22 μm thickness, larger than t_EC_ do not collapse after lifting from water (Fig. 2h and Supplementary Fig. 105). The actual thickness of PI film used to fabricate fan is 16 µm. Additionally, when assuming the barbs are cylindrical, the energy equation can be expressed as a quartic equation in terms of the radius r, which is the radius of cylinder:

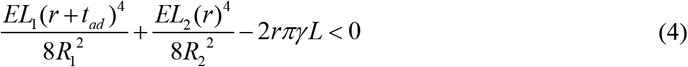

In this case, calculations using the given parameters indicate that r must be designed to be less than 11.1 µm. The extent to which different lamella and cylinder-shaped barbs deform under drag can be examined using a simplified large beam deflection model^45^. For the flexible beam loaded with a distributed load w_0_ due to leg swing as shown in Fig. 2j, nonlinear integral differential equation is given as follows:

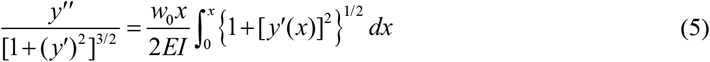

To simplify the equation we assumed *x*_0_ = *x* + Δ*x* / (*L* − Δ), where Δ is the horizontal displacement at the free end of the member (Fig. 2j), then applied boundary condition *y*′ = 0 at *x* = *L* − Δ, the nonlinear integral differential equation is simplified as follows:

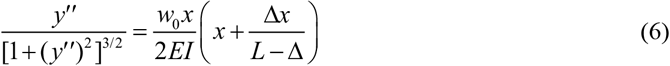

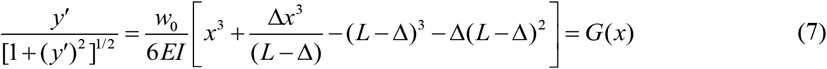

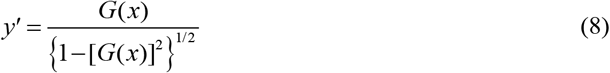

To determine *y*′(*x*), we use the Simpson’s One-Third rule to first calculate the value of the horizontal displacement Δ of the free end of the beam. This can be determined by using parameters in Supplementary Table 2 and the equation:

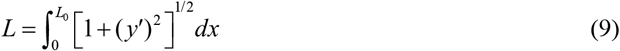

The value of Δ satisfying the length of the barb (L) is determined in four cases. The first case is a cylinder with a radius of 11.1 µm as previously calculated for collapsibility of barb, and the remaining three cases are lamella with widths of 0.03 mm, 0.1 mm, and 0.3 mm and same thickness of 16 µm. Then, we use a pseudo-linear system to determine *M*_*e*_ ′ :

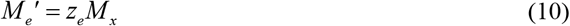

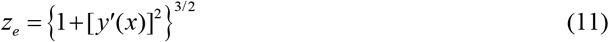

The moment-area method with the approximated *M*_*e*_ ′ is applied to determine deflection at any points in the four cases as shown in Fig. 2j (0 < *x* < *L* − Δ).

### Feedforward control for maneuvering the robot

To optimize the rapid actuation of the robot’s shape memory alloy (SMA) actuators a voltage was applied as shown in Fig. 3c. Initially, a high voltage, V1 (3 V), is applied to the actuators for 50 ms to enable the SMA to execute a full stroke rapidly. Subsequently, a reduced voltage, V2 (0.75 V), is applied to maintain the full stroke position without causing SMA overheating. Actuators R1 and L1 are connected to the right and left swing input, respectively, while R2 and L2 are connected to right and left lift input, allowing the actuator to move in a horizontal direction for swing inputs and in a vertical direction for lift inputs (Supplementary Fig. 9a). Each driving leg passes through holes in both swing and lift input, enabling it to horizontally swing or vertically lift in response to the movement of the inputs. A film guide attached to the driving leg and the frame prevents unwanted rolling of the leg. From the perspective of the right side of the robot the actuators (R1 and R2) located on the right side can position the legs in configurations ①, ②, ③, and ④ (Fig. 4B). By actuating the actuators in the desired sequence, various leg motions such as the power stroke (①→②), recovery stroke (③→④), reverse stroke (②→①), liftoff (①→③, ②→④) and touchdown (③→①, ④→②) are achievable. The coordinated actuation of four actuators allows the robot to perform a sequence of three distinct motions: advancing forward, rotating while advancing, and rotating in place (Supplementary Fig. 9b).

The robot’s rotating motion comes in two types. The first is rotation while advancing. The process begins with the actuation of R2 and L2 to advance the robot forward with a power stroke (①→②). By actuating only R2 while maintaining the operation of R1 and L1, the right leg is lifted above the water (②→③) with the other left submerged (②) inducing the robot to rotate around the submerged fan. This motion results from the reaction drag acting on the submerged fan, which is opposite to moving direction of the robot. The lifted right leg is returned to its initial state through a recovery stroke (③→④→①), while the submerged left leg is kept in place for the next motion (②). The second method of rotation, rotation in place, follows rotation while advancing. Here, the left leg remains in state ② and the other in state ①; swinging each leg in opposite direction allows for rapid on-the-spot rotation. The left leg performs a reverse stroke (②→①) by deactivating L1, while the other performs a power stroke (①→②) by activating R1, enabling rapid rotation in place. The first step of forward motion is actuating R1 and L1, causing both legs to perform a power stroke (①→②). By maintaining the operation of R1 and L1 while activating R2 and L2, the legs are lifted above the water (②→③). Deactivating the actuators in sequence returns the legs to their initial state through a recovery stroke (③→④→①).

### Model of energy balance for a rhagovelia fan

The surface energy and bending energy for rhagovelia fan’s barb is determined using Equation (1) and (2). The Young modulus of the barb is measured using AFM Supplementary Table 3 provides the parameters for calculating the surface energy and bending energy of the Rhagovelia fan. The surface area (A_robot_∼3 mm^2^) of the individual robot fan is 1000 times higher than the individual Rhagovelia fan barbs (A_rhag_∼0.0035 mm^2^), suggesting that higher dimensions enhance the more surface energy in robots compare to the Rhagovelia. The average radius of curvature is listed in the Supplementary Table 4 for individual barbs, and the corresponding map across the length, with color representing the curvature value for individual barbs. The surface energy magnitude is *O* (10^−3^ µJ), which is 1000 times higher of the bending energy (E_b_) magnitude *O* (10^−6^ µJ). The surface energy (E_s_) is higher than the bending energy (E_b_) for a single barb, suggesting that the Rhagovelia fan prioritizes surface interactions over structural flexibility (Supplementary Fig. 2e).

### Theoretical opening time for barb of Rhagovelia and robot fan

In the fixed boundary system, capillary rise is dominated by surface tension and viscous forces^46^. We observed that the liquid rise in the Rhagovelia fan depends on the movement and deformation of individual barbs. For deformable boundary system, the capillary rise equation is influenced by viscous forces, surface tension forces, and elastic forces (due to flexible fibers).

For the flexible long fan, the dynamics of its opening are influenced by the liquid rising into the barb. We calculate the theoretical opening time (τ) using the following equation^47^.

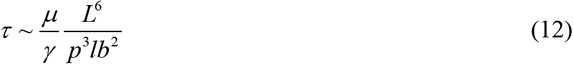

Here, *lb* is the capillary bending length^48^, *p* is pore width from top, *L* is the barb length, *μ* is viscosity, and *γ* is surface tension.

For a cylinder

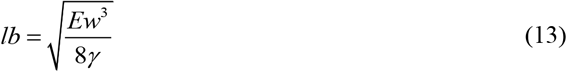

For a robotics fan, where the thickness is much smaller than the width

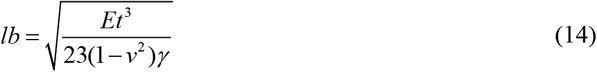

Our finding indicates that the fan opening time is dependent on the liquid’s rise and the opening of the fan, where the channel width continuously changes with the rising of liquid. This equation demonstrates that faster capillary rising results in greater spreading.

The opening time (τ) is influenced by several parameters, including the length of the barbs, their flexibility, the width of the barbs, the properties of the water, and the dimension of the pores or channels (both length and width) as listed in Supplementary Table 1. The channel width is determined by the opening of the fan (Supplementary Fig. 2f). The fan consists of multiple barbs, with two barbs forming a single channel, and the total number of channels is (n-1). We measured the maximum opening width

(p) of individual channels for both the fan and the robot (Supplementary Fig. 2f), which ranges from 40 µm to 200 µm for the insect fan and from 0.5 mm to 1.5 mm for the robot. Furthermore, we calculated the theoretical opening time for the maximum open channel width (Supplementary Fig. 2g). This theoretical time is of a similar order of magnitude, *O*(∼10 ms) and *O*(∼300 ms), as the experimental opening times observed in both cases.

### Theoretical opening time for the barbules of the Rhagovelia and robot fans

We measured the pore width between the individual two barbules (Supplementary Fig. 2b) in the SEM image of a single barb. The parameters used for the calculation are listed in Supplementary Table 5. Confocal images of the barb and barbules of the Rhagovelia fan suggest that both barb and barbules are similar materials, as the fan emits similar color using autofluorescence (Supplementary Fig. 2k and 2l). When the sample is excited with a laser wavelength of 488 nm. We consider the young modulus of the barbules to be same as that of the barb. The barbules pore width varies from 5 to 25 µm (Supplementary Fig. 2i), with corresponding theoretical opening times on the order of O (∼1ms), which aligns with the range of experimental observations (Supplementary Fig. 2h).

### Rhagovelia barb modulus measurements

We determined the young modulus of the Rhagovelia fan’s barb through a series of steps. Initially, we isolated the fan from the organism and spread it on the glass slide surface to measure the young modulus (Supplementary Fig. 6a). Subsequently, we placed the sample glass slide in the atomic force microscopy (AFM, Asylum MFP-3D Bio) setup (Supplementary Fig. 6b) and focused placed the AFM tip on the one of the barbs (inset of Supplementary Fig. 8d). We used the 5 µm diameter polystyrene sphere attached to the AFM probe cantilever to perform the indentation test. We selected the 7×7 µm^2^ area map to quantify the young modulus of the fan (Supplementary Fig. 6c). Our measurements yielded an average modulus of approximately 15 MPa for barb, as represented in the histogram (Supplementary Fig. 6d).

