## Supplementary Information for "Ultrafast elastocapillary fans control agile maneuvering in ripple bugs and robots"

### Table of contents:

- **Supplementary Figures**


- Supplementary Figure 1. Opening and closing of Rhagovelia fan.
- Supplementary Figure 2. Structural analysis of Rhagovelia fan barbs and barbules.
- Supplementary Figure 3. Fan collapse.
- Supplementary Figure 4. Fabrication of artificial fan.
- Supplementary Figure 5. Bending and surface energy model of artificial fan.
- Supplementary Figure 6. Analysis of Young modulus of the Rhagovelia fan.
- Supplementary Figure 7. Velocity Field.
- Supplementary Figure 8. Intact and disabled fan.
- Supplementary Figure 9. Control of actuating leg and rhagobot motions.
- Supplementary Figure 10. Vorticity field using PIV.
- Supplementary Figure 11. SEM image of a barb and barbules.
- Supplementary Figure 12. DeepLabCut tracking.


- **Supplementary Tables**


- Supplementary Table 1. Parameters for calculating the opening time of the Rhagovelia fan and the robot fan.
- Supplementary Table 2. Parameters for energy model and large beam deflection.
- Supplementary Table 3. Parameters for the calculation of bending and surface energy of the Rhagovelia fan.
- Supplementary Table 4. Average radius of curvature for individual Rhagovelia barbs.
- Supplementary Table 5. Parameters for calculating the opening time of the Rhagovelia fan barbules.
- Supplementary Table 6. Two-way Repeated measures ANOVA.
- Supplementary Table 7. Semi-aquatic robot data.


- **Description of Supplementary Videos**


- Supplementary Video 1. Rhagovelia’s fan and rowing performance
- Supplementary Video 2. Engineered fan and agile interfacial locomotion of a robot


- **Description of Source data**


- Source data Fig. 1. Fan spreading
- Source data Fig. 2. Fan energy and large beam deflection modelling
- Source data Fig. 3. Rhagovelia turning kinematics
- Source data Fig. 3. Disable and intact Fan experiment
- Source data Fig. 4. Rhagobot performance with and without fan

**Supplementary Figures**

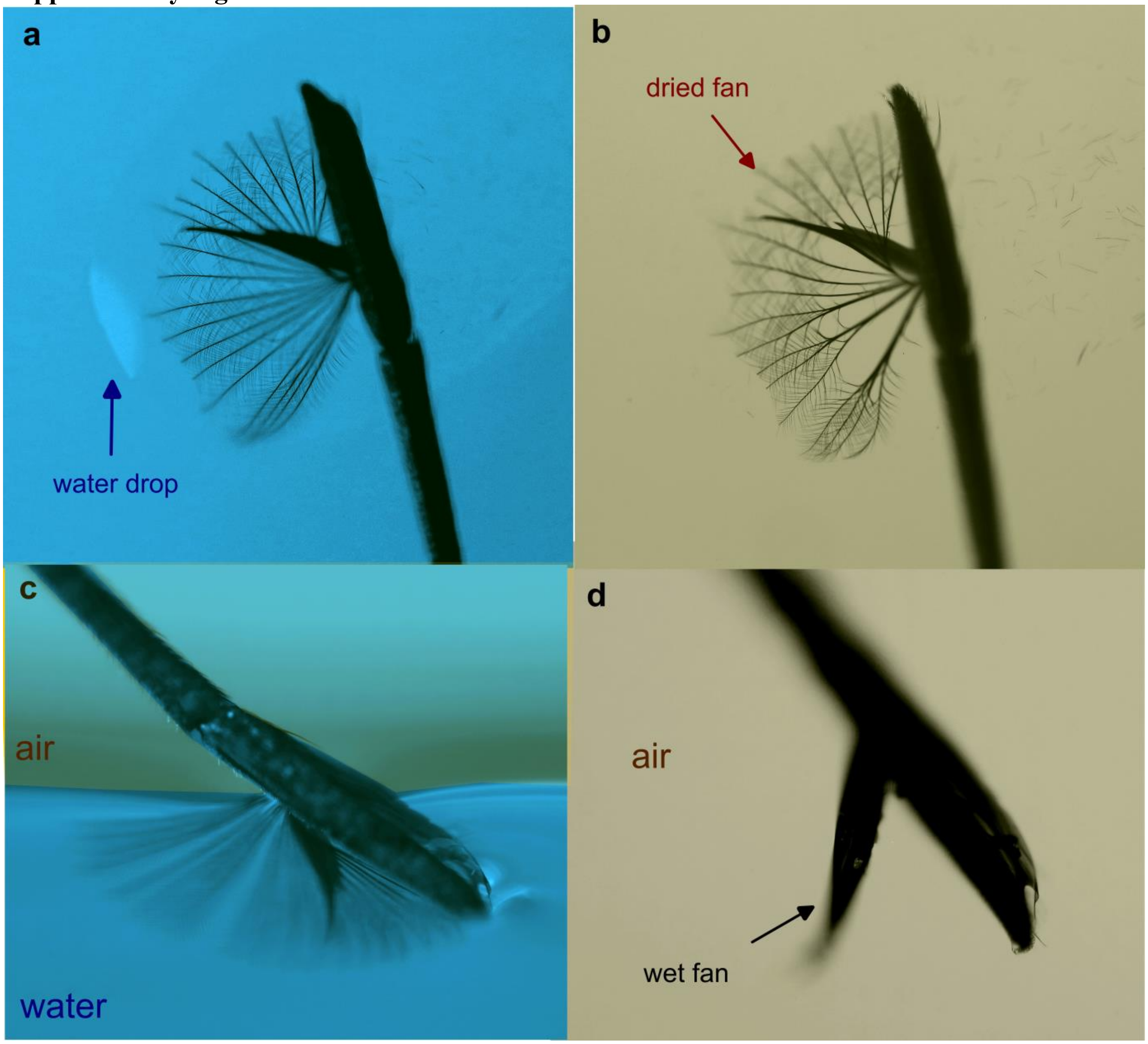


**Figure S1**. **Opening and closing of Rhagovelia fan**. **a,** Open fan in a water droplet. **b,** Dried fan remains fully spreaded. **c,** Isolated Leg and fan. If the claw is open It spreads when submerged underwater. **d,** Fan collapses towards the claw when removed from water.

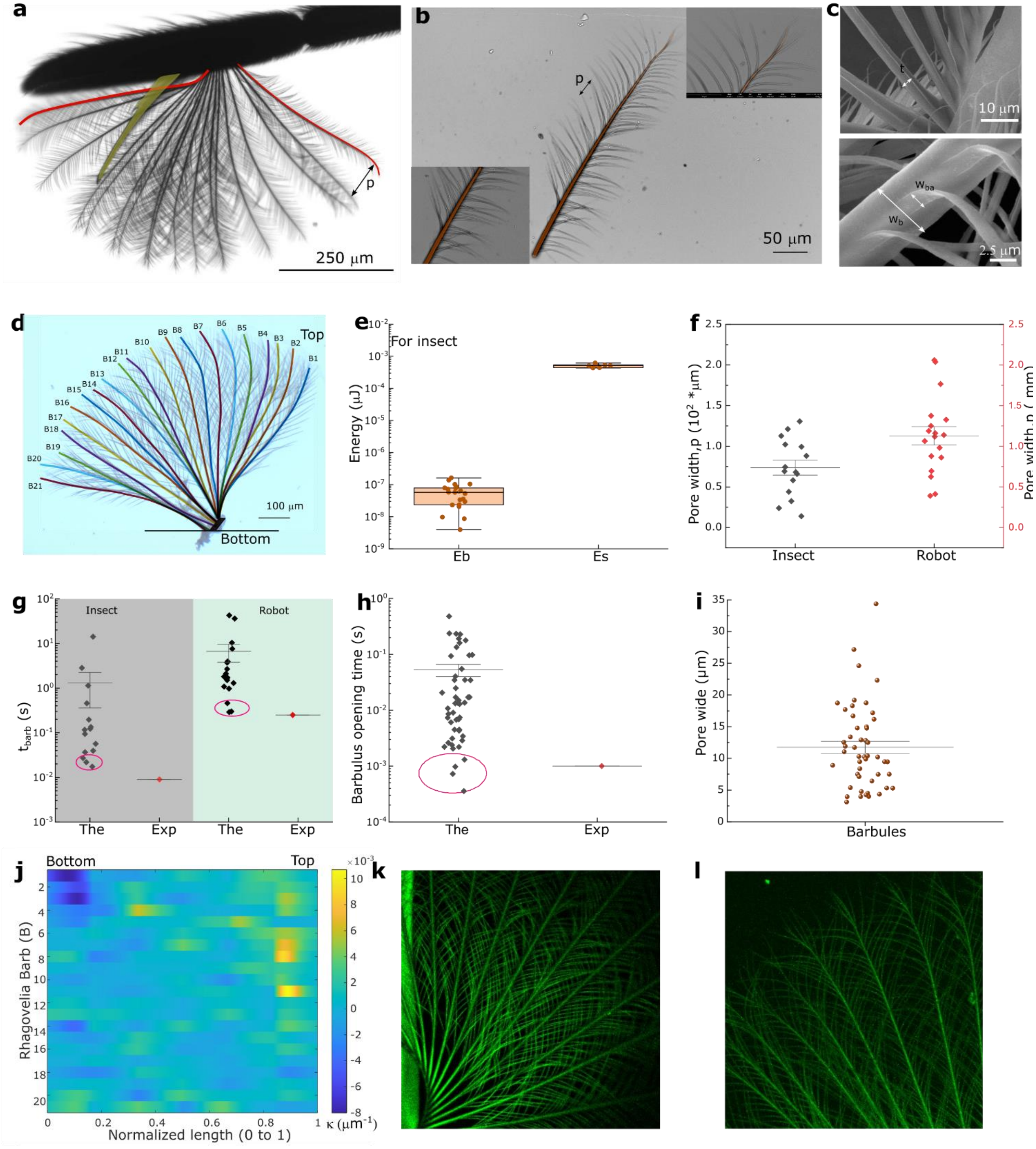


**Figure S2. Structural analysis of Rhagovelia fan barbs and barbules. a,** Rhagovelia fan image defining the parameters: pore width (p), curve length highlighted in red. **b,** Scanning electron microscopy (SEM) image of a single barb with barbules, illustrating the barb's thickness being greater at the bottom and tapering towards the top. **c,** SEM image displaying the barb width ($w_b$) and thickness (t), as well as the barbule width ($w_{ba}$) and thickness ($t_{ba}$) of the Rhagovelia fan. **d,** X and Y coordinates for an open barb (for 21 barb) of the Rhagovelia fan. **e,** The plot illustrates the bending energy ($E_b$) for the curvature of individual fan barbs and the surface energy ($E_s$) for the width ($w_b$) individual fan barbs in the Rhagovelia fan. **f,** The scatter interval plot displays the fan pore widths (p) at the top measurements for insects and robots. **g,** The scatter interval plot for opening time ($t_{barb}$) compares the theoretical and experimental measurements of all pore widths (p) of the barb in both insect and robotic systems. **(h-i),** Theoretical and experimental values of barbules opening time ($t_{barbules}$) for different pore widths (p). **j,** Plot of individual barb curvature ($\kappa$) across the normalized length. **(k-l)**, Confocal autofluorescence image of the Rhagovelia fan at 20x magnification, from 488 nm excitation.

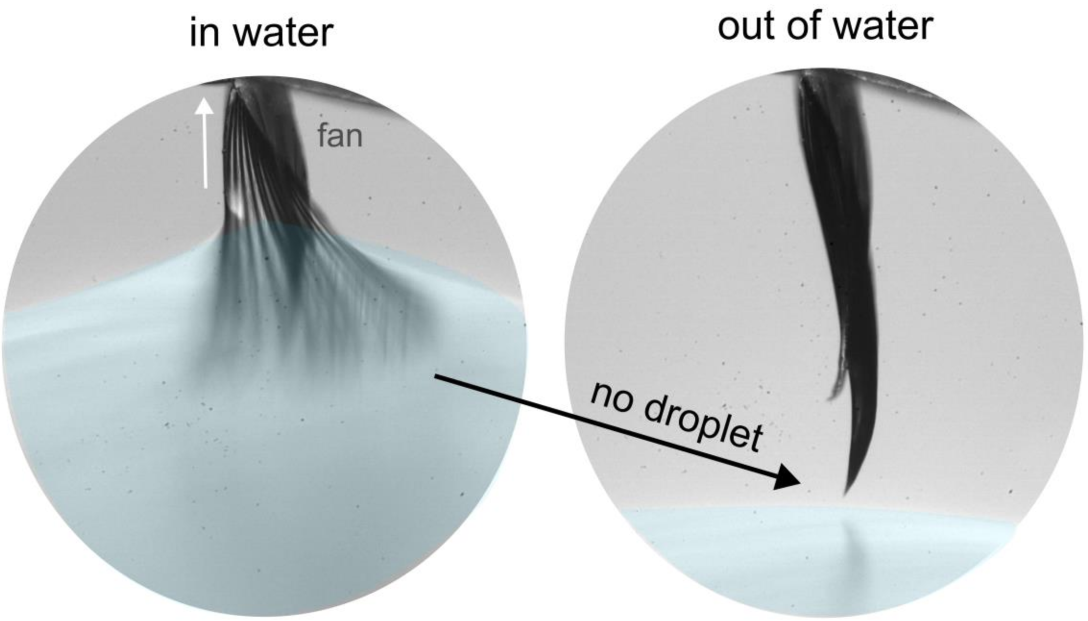


**Figure S3. Fan collapse**. Fan inside the water fully spreads, but when pulled out of the water it collapses. Notice that the collapsed fan forms a sharp tip and it does not retain a water droplet.

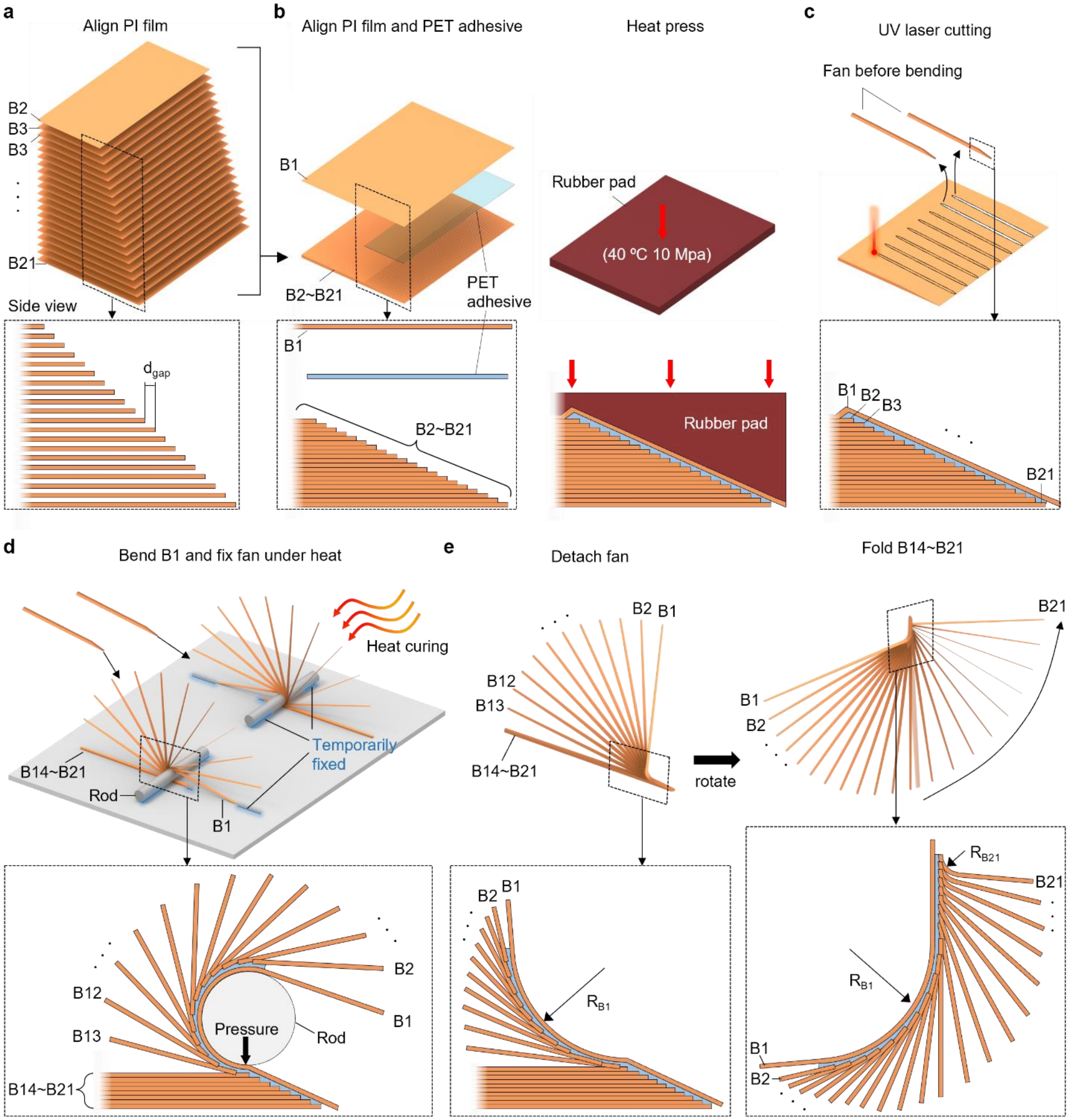


**Figure S4. Fabrication of artificial fan. a,** Twenty sheets of pi film layers (B2~B21) are aligned with a length difference of $d_{gap}$. **b and c,** To attach the B1 layer to the other 20 layers (B2~B21), a PET adhesive is placed in between and pressed with a rubber pad. **d,** The fan in its pre-unfolded state is released through UV laser machining. **e,** The B1 layer is fixed and wrapped around a rod to unfold the fan, and heat is applied. **f,** The fan is released from its fixation, and the remaining B14 to B21 layers are folded to form the shape of the fan.

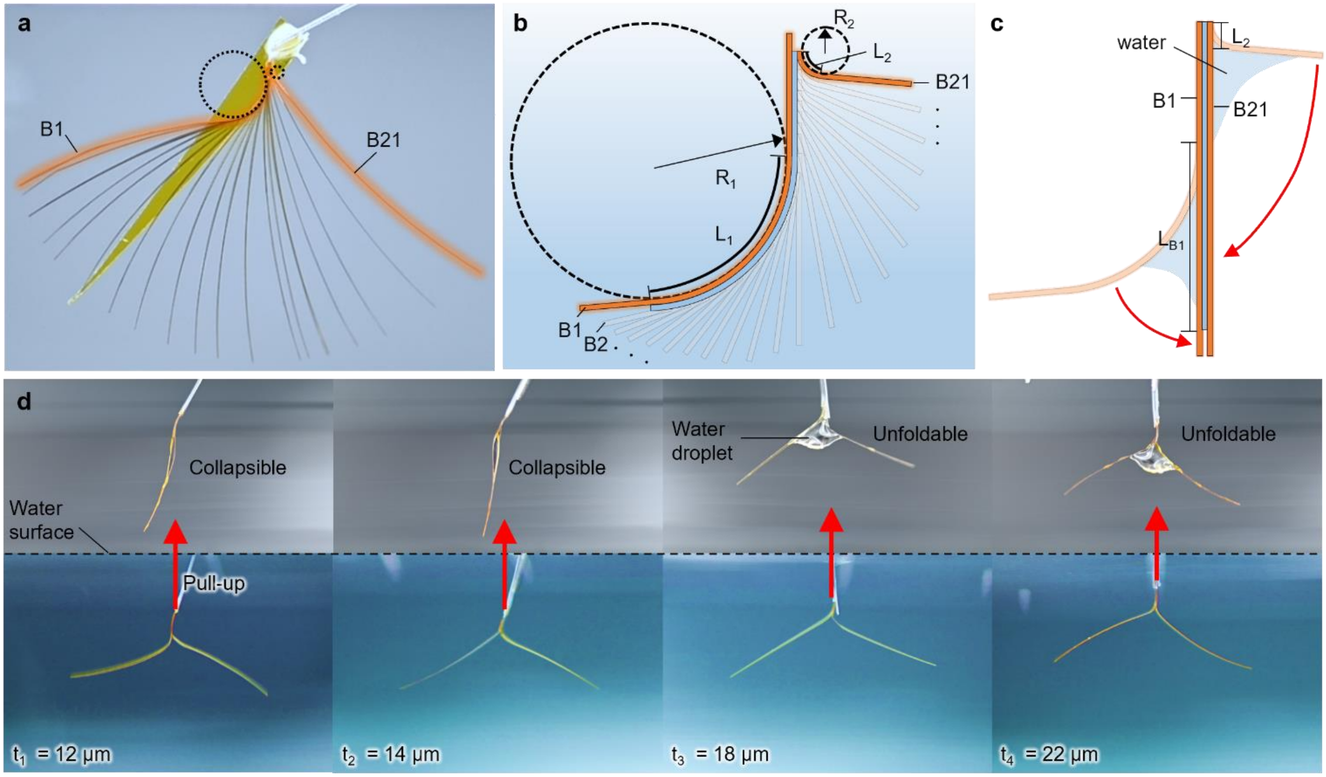


**Figure S5. Bending and surface energy model of artificial fan. a,** Artificial fan is fully spread under water with the most bent barbs at both ends Highlighted. Each dotted circle is in contact with the bent part of the barb. **b,** A zoomed-in illustration of the state of spread fan shown in **a**. Bending radius (R), bent length (L) to calculate bending energy are presented for each barb. **c,** The fan is in collapsed state due to capillary force when removed from the water. **d, e, f and g**, the 12, 14, 18 and 22 um thickness of fan are lifted from the water to verify minimum thickness for collapsibility.

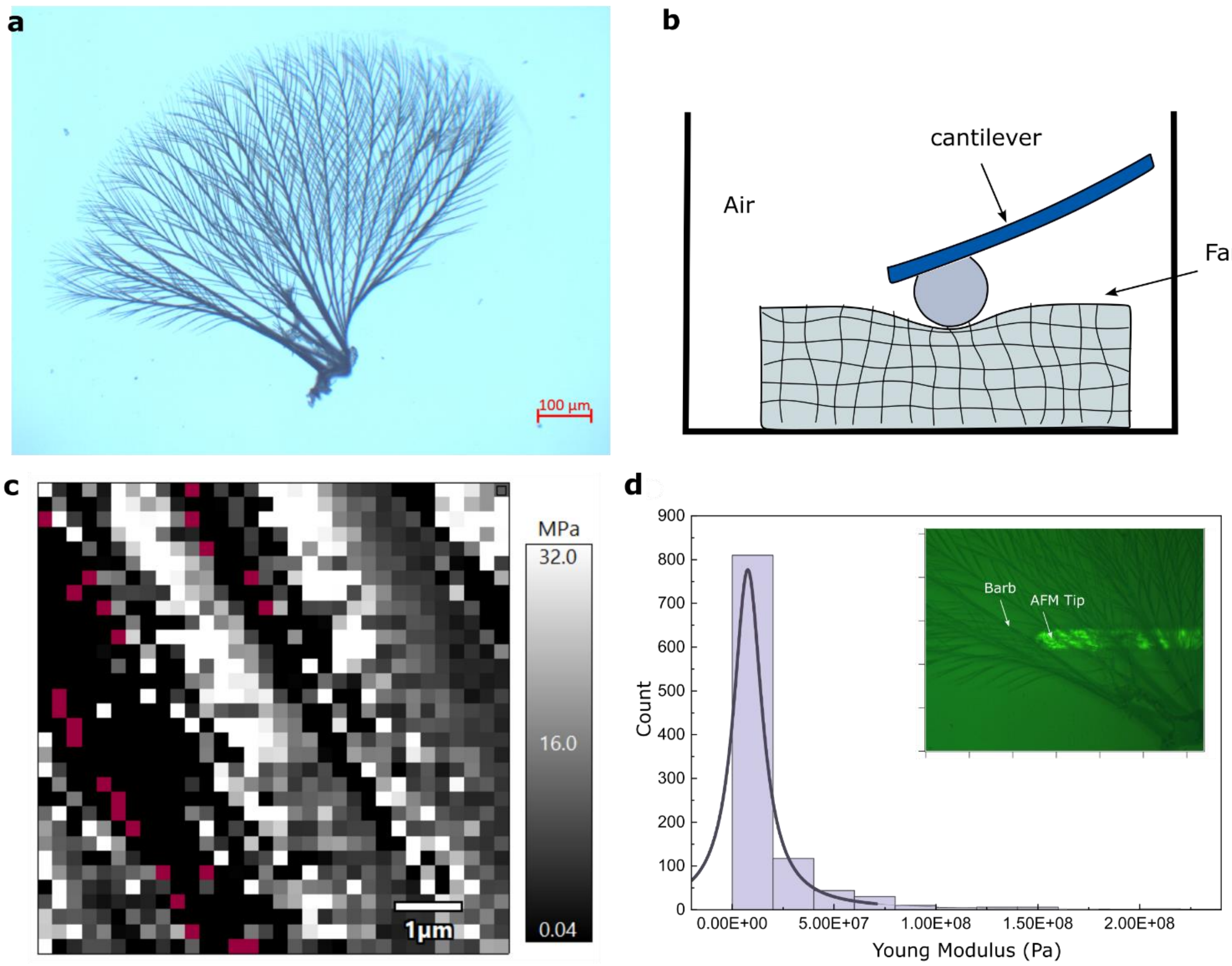


**Figure S6. Analysis of Young modulus of the Rhagovelia fan. a,** Optical microscopy image of the Rhagovelia fan. **b,** Setup of the AFM. **c,** Young modulus map of the 7x7 µm$^2$ area. **d,** Histogram plot of the young modulus of the barb with an inset image showing the AFM tip on the fan barb.

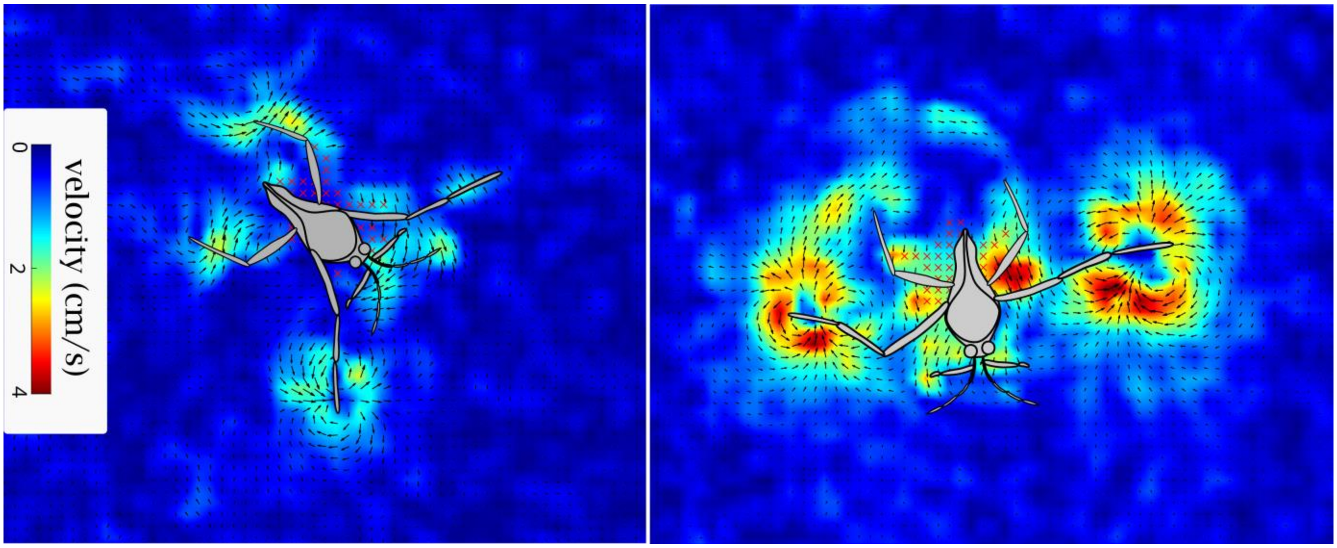


**Figure S7. Velocity Field**. Rhagovelia during turning (right) and during the power stroke (left).

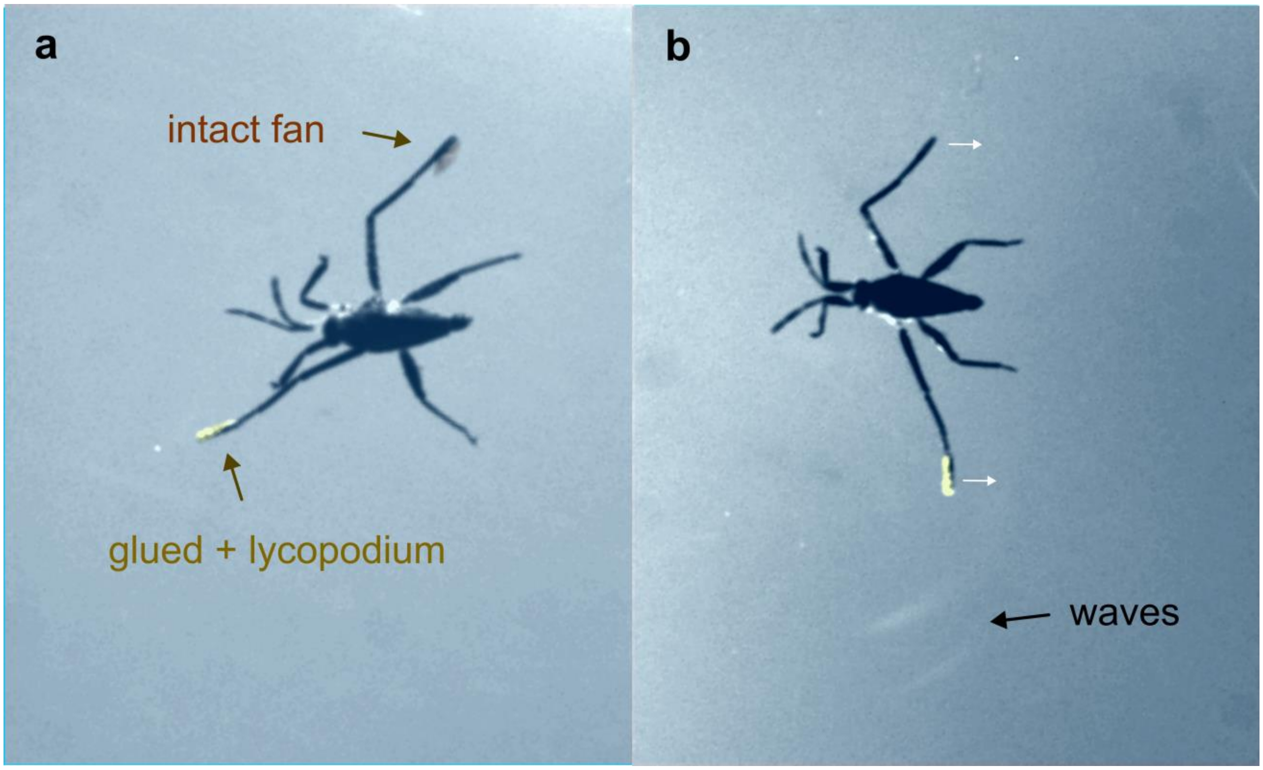


**Figure S8. Intact and disabled fan**. **a,** Individual showing a right leg with a functional fan and a left leg with a disabled fan, which was glued and dusted with hydrophobic particles (lycopodium). **b,** Individual showing capillary waves produced during the propulsive stroke by the leg with a disabled fan.

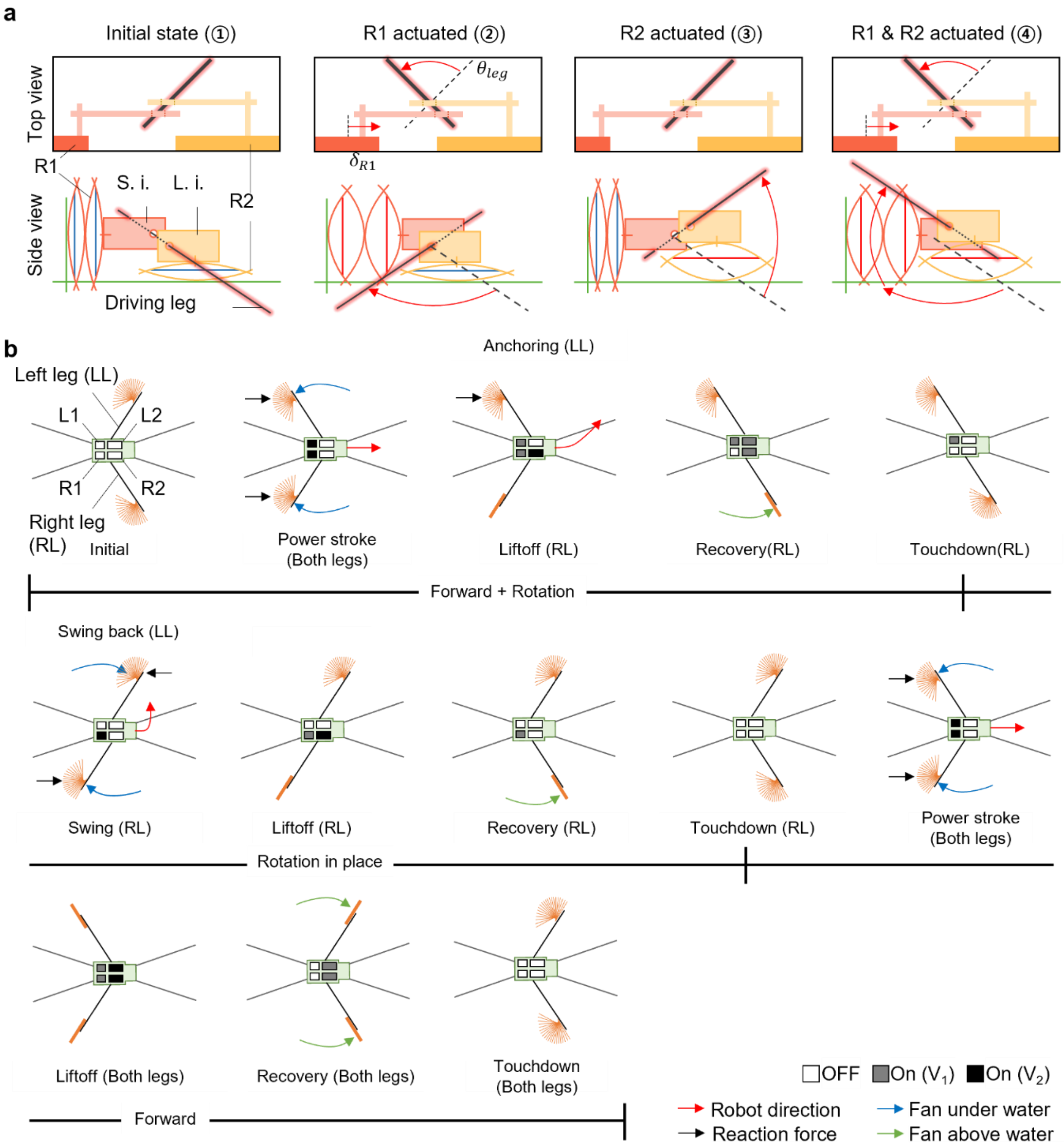


**Figure S9. Control of actuating leg and rhagobot motions. a,** The exploded top and side views of the robot's right side. Linear actuation of R1 and R2 are converted to leg' rotational stroke via the swing and lift inputs. Stroke of R1 ($\delta_{R1}$) and R2 ($\delta_{R2}$) induces 80° and 40° of leg rotation ($\theta_{leg}$) in swing (②) and lift (③) directions respectively. **b,** The top view of the robot showing robot's moving direction according to leg movement induced by actuators.

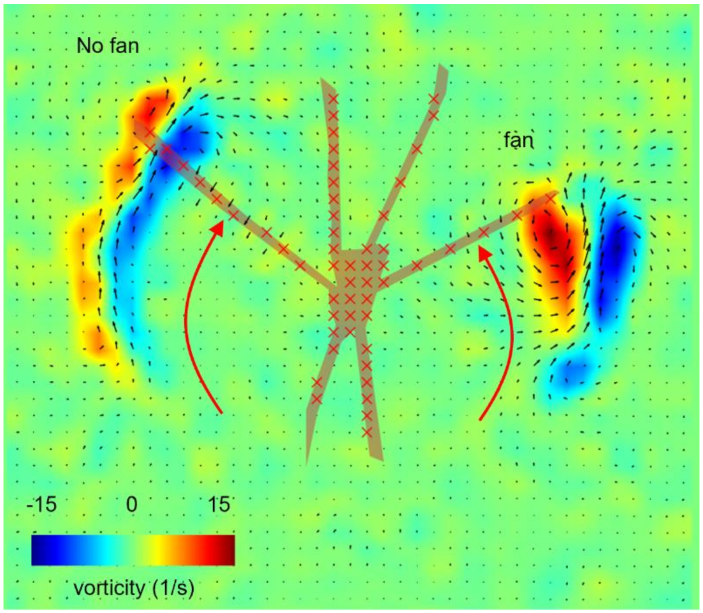

**Figure S10. Vorticity field using PIV.** Comparison of a leg with a fan and a superhydrophobic coated leg withouta fan.

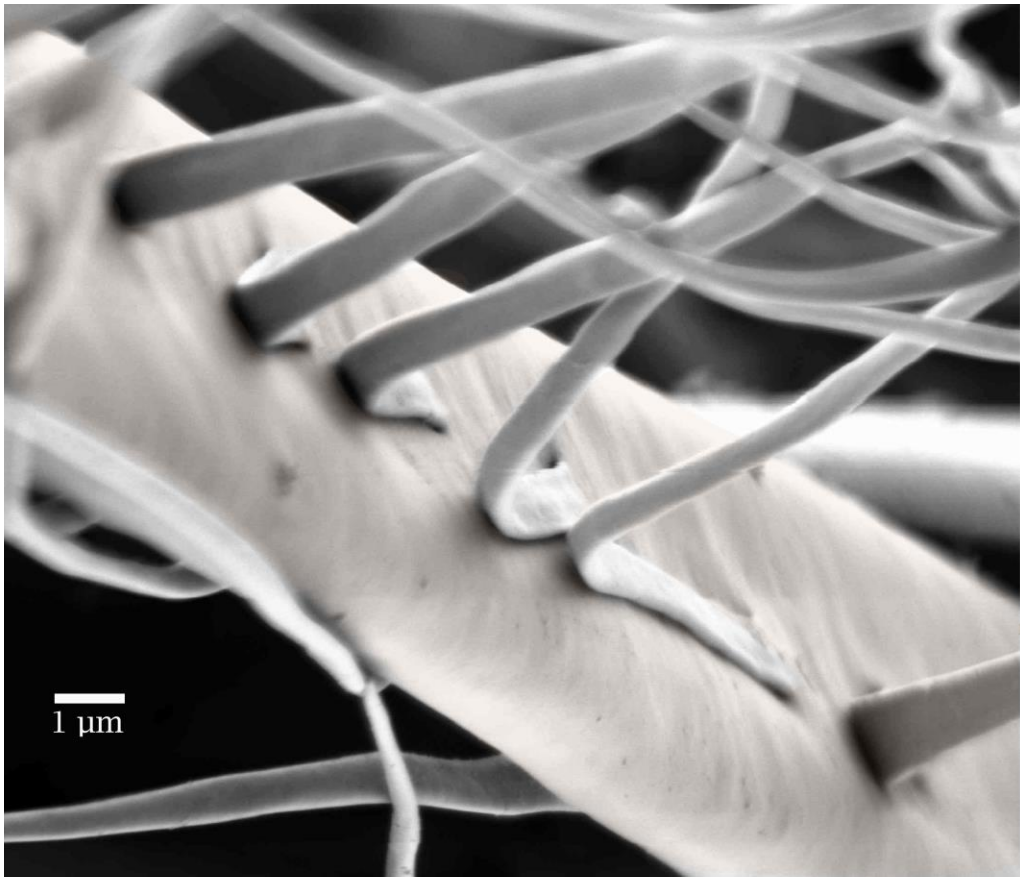


**Figure S11. SEM image of a barb and barbules**. Notice that that the barb is slightly depressed in its middle part.

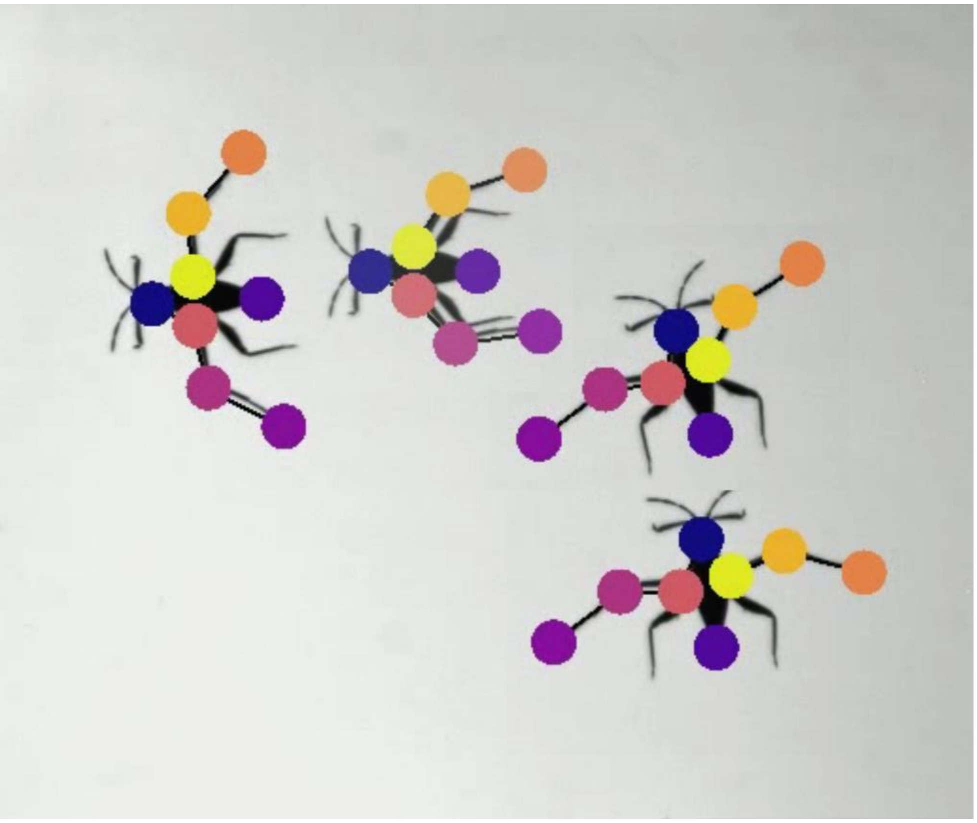

**Figure S12**. **DeepLabCut tracking.** Example of merged video frames showing automatic tracking of a Rhagovelia individual during turning.

#### Supplementary Tables

| Parameter | Symbol | Value (Robotic fan) | Value (Insect fan) | Unit |
|---|---|---|---|---|
| Water viscosity | m | 0.001 | | Pa·s |
| Surface tension | g | 72.8 | | mN |
| Young modulus of Barb | E | 2000 | 15 | MPa |
| Barb length | L | 0.01 | 5.87E-04 | m |
| Barb thickness | t | 16E-06 | 2.37E-06 | m |
| Barb width | $w_b$ | 3.00E-04 | 6.00E-06 | m |
| Poisson ratio | n | 0.37 | 0.5 | - |
| Experimental opening time | $t_{exp}$ | ~250 | ~9 | ms |
| Theoretical opening time | $t_{the}$ | ~O(300) | O(~10) | ms |
| Pore width (maximum) | p | ~2 | ~0.10 | mm |

**Table S1. Parameters for calculating the opening time of the Rhagovelia fan and the robot fan.**

| Parameter | Symbol | Value | |
|---|---|---|---|
| Surface tension | $\gamma$ | 72.8 | mN/m |
| Young's Modulus of polyimide film | $E$ | 2 | Gpa |
| Barb length | $L$ | 10 | mm |
| Curved length of B1 and B21 | $L_1$, $L_2$ | 2.03, 0.133 | mm |
| Bending radius of B1 and B21 | $R_1$, $R_2$ | 1.418, 0.218 | mm |
| Barb Width | $w_1$, $w_2$, $w_L$ | 0.03, 0.1, 0.3 | mm |
| Adhesive thickness | $t_{ad}$ | 1 | μm |
| Distributed force acting on the barb | $w_0$ | 6.3E-7 | N/mm |
| Barb thickness | $t$ | 16 | μm |
| Horizontal displacement | $\Delta 1$, $\Delta 2$, $\Delta L$, $\Delta r$ | 3, 0.005, 0, 5.3 | mm |

**Table S2. Parameters for energy model and large beam deflection**

| Parameters | Symbol | Value | Unit |
|---|---|---|---|
| Young modulus of Barb | E | 1.50E+07 | MPa |
| Barb length | $L_b$ | 587±0.012 | mm |
| Barb thickness | T | 2.37±0.24 | mm |
| Barb width | $w_b$ | 6.00±0.52 | mm |
| Number of Barb | N | 21 | |
| Barbules length | $L_{ba}$ | 70±7.97 | mm |
| Barbules thickness | $t_{ba}$ | 0.33±0.075 | mm |
| Barbules width | $W_{ba}$ | 1.63±0.13 | mm |
| Single barb surface Energy | $E_s$ | 5.13E-10 | J |
| Radius of curvature | $R_b$ | 515 | mm |
| Single barb bending Energy | $E_b$ | 1.11E-13 | J |

**Table S3. Parameters for the calculation of bending and surface energy of the Rhagovelia fan.**

| Barb | Average radius of curvature (μm) |
|---|---|
| B1 | 460.1933 |
| B2 | 535.6186 |
| B3 | 425.7131 |
| B4 | 530.2227 |
| B5 | 609.3845 |
| B6 | 676.59 |
| B7 | 586.1665 |
| B8 | 637.3486 |
| B9 | 1116.071 |
| B10 | 998.004 |
| B11 | 710.7321 |
| B12 | 1739.13 |
| B13 | 909.0909 |
| B14 | 745.1565 |
| B15 | 1129.944 |
| B16 | 1189.061 |
| B17 | 2732.24 |
| B18 | 934.5794 |
| B19 | 1848.429 |
| B20 | 713.7759 |
| B21 | 615.0062 |

**Table S4. Average radius of curvature for individual Rhagovelia barbs.**

| Parameter | Symbol | Value | Unit |
|---|---|---|---|
| Young modulus of Barbules | E | 15 | MPa |
| Barbules length | L | 70E-06 | m |
| Barbules thickness | t | 0.33E-06 | m |
| Barbules width | $w_b$ | 1.63E-06 | m |
| Poisson ratio | N | 0.5 | |
| Experimental opening time | $t_{exp}$ | ~1 | ms |
| Theoretical opening time | $t_{the}$ | O(~1) | ms |
| Pore width (maximum) | p | ~20 | μm |

**Table S5. Parameters for calculating the opening time of the Rhagovelia fan barbules.**

| Term | Sumsq | df | Error | F value | P value |
| --- | --- | --- | --- | --- | --- |
| Fan (active/inactive) | 0.1086 | 3 | 0.0068100 | 47.8351 | 0.006196 ** |
| Leg (left/right) | 0.0029 | 3 | 0.0054850 | 1.5639 | 0.299754 |

**Table S6. Two-way Repeated measures ANOVA.** We to compare the effect of fan (intact vs disabled) and leg treated (left or right) in the maximal speed reached by the leg tip during the power stroke. Test was performed in R V4.4.1.

| Source | Mass (g) | Relative speed (BL/s) | Turning speed (°/s) | Propulsion mechanism | Tethered/ Untethered | Actuator |
|---|---|---|---|---|---|---|
| [12] | 3.9 | 1.33 | 23 | Hydrophobic leg | Tethered | DC motor |
| [17] | 3.88 | 1 | - | Hydrophobic leg | Tethered | DC motor |
| [18] | 21.75 | 0.27 | 12 | Hydrophobic leg | Untethered | DC motor |
| [13] | 7.5 | 0.15 | - | Hydrophobic leg | Tethered | DC motor |
| [14] | 1 | 0.3 | 28.65 | Hydrophobic leg | Tethered | Piezoelectric |
| [19] | 6.13 | 0.58 | 45.8 | Hydrophobic leg | Untethered | DC motor |
| [15] | 4.9 | 0.71 | 24.8 | Hydrophobic leg | Tethered | DC motor |
| [8] | 510 | 0.39 | 22.5 | Rigid pad | Untethered | DC motor |
| [9] | 863 | 1.4 | 45 | Rigid pad | Untethered | DC motor |
| [10] | 208 | 0.69 | - | Rigid pad | Tethered | DC motor |
| [7] | 137 | 0.62 | 35 | Rigid pad | Tethered | DC motor |
| [11] | 439 | 0.74 | - | Rigid pad | Untethered | DC motor |
| [22] | 1.6 | 0.7 | 32.7 | Passive flap | Tethered | Piezoelectric |
| [23] | 0.056 | 0.28 | 8.25 | Passive flap | Tethered | Shape memory alloy |
| [24] | 2.51 | 0.22 | 15.43 | Passive flap | Untethered | Electroosmotic hydrogel |
| [25] | 14.3 | 0.77 | 20.45 | Passive flap | Tethered | DC motor |
| [26] | 22.65 | 1.23 | - | Passive flap | Tethered | DC motor |
| **This work** | **0.24** | **1.47** | **87** | **Collapsible fan** | **Tethered** | **Shape memory alloy** |

**Table S7. Semi-aquatic robot data.**

#### Description of Supplementary Videos

**Supplementary Video 1. Rhagovelia's fan and rowing performance**

[00:06] *Rhagovelia obesa* showing fans.

[00:10] SEM image of the fan highlighting flat barbs and barbules.

[00:10] Isolated fan placed and removed from a water droplet.

[00:25] Fan collapses after removing it from the water.

[00:29] Rhagovelia's fan actuation during rowing.

[00:38] Rhagovelia's fan actuation during swimming.

[00:47] Rhagovelia using an asymmetric reverse stroke for turning.

[00:57] Rhagovelia using a symmetric reverse stroke for braking.

[01:10] Rhagovelia's rowing performance using Deep Lab Cut.

[01:20] Vortical wakes produced by Rhagovelia (top view).

[01:45] Fan's vortical wake (side view).

[01:56] Capillary waves produced by Rhagovelia (top view).

[02:04] Rhagovelia rowing performance with intact and disabled fans.

[02:15] Wake differences of an individual while rowing with intact and disabled fans.

[02:45] Rhagovelia in natural streams.

[03:00] Particles used to perform PIV in natural streams.

[03:11] Day and night activity of Rhagovelia (from August 2023 to April 2024).

**Supplementary Video 2. Engineered fan and agile interfacial locomotion of a robot**

[00:06] Rhagovelia-inspired robot, "rhagobot," equipped with a robotic fan

[00:12] Robotic fan comprising a claw and 21 barbs

[00:19] Microscopic image of the robotic fan and SEM image of biological fan both highlighting flat barbs.

[00:24] Robotic fan passively spreading and collapsing, driven by elsastocapillarity.

[00:34] High-speed video of the electrocapillary morphing fan.

[00:56] Rhagobot swinging its driving leg with the fan attached at the end (side view).

[01:11] Robotic fan retaining its shape underwater while striking.

[01:25] Comparison of the robots maneuvering with and without the fan (top view).

[01:35] Comparison of the robots advancing and braking with and without the fan.

[01:50] PIV showing vorticity differences in robots with and without the fan.

#### Description of Supplementary Videos

##### Source data Fig. 1. Fan spreading

Raw data shows the time series of spreading fan angle. Angles are in degrees. First column in each sheet represent time (s). Sheets 1 and 2 are data of isolated fan spreading angles during opening and closing, respectively. Sheets 3 and 4 are data of fan spreading angles during opening and closing of individuals during rowing, respectively. Sheets 5 to 8 are data of fan and claw spreading angles during opening and closing of individuals during swimming, respectively.

##### Source data Fig. 2. Fan energy and large beam deflection modelling

Raw data shows bending and surface energy modelling for a flat beam (rectangular cross-section) and a cylinder beam (sheet 1), and large beam deflection for three cases of flat beams and the cylinder beam (sheet 2). Sheet 3 and 4 contain the time series of the fan opening and closing angle of four trials, respectively.

##### Source data Fig. 3. Rhagovelia turning kinematics

Raw data shows the time series of the body angle (sheet 1), left-leg angle (sheet 2) and right-leg angle (sheet 3). Each column (Column 2- Column 160) represents different trials. Data is in degrees.

##### Source data Fig. 3. Disable and intact Fan experiment

Raw data shows the time series of the left (sheet 1) and right (sheet 2) leg tip speed of 4 individuals (a,b,c,d) during control, one fan disabled, and two fans disabled. Sheet 3 contains the maximal tip-leg speed for each treatment per individual.

##### Source data Fig. 4. Rhagobot performance with and without fan

Raw data shows trajectory of a robot with and without fan (sheet 1). Sheet 2 contains travel distance of two robots advancing for six trials, respectively. Sheet 3 contains travel distance and turning angle of two robots rotating while advancing for 8 trials, respectively. Sheet 4 contains turning angle of two robots rotating in place for 8 trials.
